# Glutamate is a key nutrient for *Porphyromonas gingivalis* growth and survival during intracellular autophagic life under nutritionally limited conditions

**DOI:** 10.1101/2024.07.08.602514

**Authors:** Nityananda Chowdhury, Bridgette Wellslager, Hwaran Lee, Jeremy L. Gilbert, Özlem Yilmaz

## Abstract

*Porphyromonas gingivalis* survives in special autophagic vacuoles that serve as major replicative habitats in human primary gingival epithelial cells (GECs). As an asaccharolytic strict anaerobe, *P. gingivalis* is dependent on amino acids and peptides for nutrient sources. However, it is largely unknown as to *P. gingivalis’* metabolic processing under the nutritionally limited intracellular environments such the vacuoles, especially the preferred amino acids and associated-metabolic machineries. Here we elucidate that a Glutamate (Glu) catabolic enzyme, glutamate dehydrogenase (GdhA) is highly enriched in the isolated *P. gingivalis*-containing vacuoles. Interestingly, we found that *P. gingivalis* induces conversion of intracellular glutamine pool to Glu determined by analyses of the *P. gingivalis-*containing vacuoles and the whole infected-GECs. Critically, exogenous Glu-Glu dipeptide, a simple precursor of Glu, significantly increases the size of isolated intact *P. gingivalis* containing-vacuoles and live wild-type *P. gingivalis* numbers in GECs. In contrast, the isogenic GdhA-deficient-strain, Δ*gdhA* displayed a significant growth defect with collapsed-vacuoles in GECs. Next, we confirmed that *P. gingivalis* uptakes ^14^C-Glu and it preferentially utilizes Glu-Glu-dipeptide using a nutritionally reduced Tryptic-Soy-Broth (TSB) media supplemented with Glu-Glu. Contrary, Δ*gdhA*-strain showed no detectable growth especially in nutritionally reduced TSB media with Glu-Glu. Using Atomic-Force-Microscopy, we observed that, wild-type *P. gingivalis* but not Δ*gdhA* strain notably increased the cell volume upon Glu-Glu supplementation, an indicator of higher metabolism and growth. Utilization of a human gingiva-mimicking organoid-system further validated the importance of Glu as an essential nutrient for the intramucosal colonization of *P. gingivalis* via the protected replicative vacuoles in GECs.

**Importance:** This study reveals that *P. gingivalis* heavily depends on preferential utilization of Glutamate (Glu) for autophagic vacuolar growth and survival in human GECs. Several novel observations are made to support this: (i) GdhA of *P. gingivalis* is highly enriched in these vacuoles, (ii) *P. gingivalis* induces a large conversion of intracellular glutamine to Glu, (iii) size of vacuoles are significantly increased in the presence of Glu-Glu in *P. gingivalis* wild-type strain infection which is opposite in a Δ*gdhA* strain, (iv) *P. gingivalis* uptakes ^14^C-Glu and preferentially utilizes Glu-Glu dipeptide, (v) similarly, wild-type strain shows growth increase in a nutritionally reduced bacterial culture media, and (vi) finally, Glu-Glu supplementation increases bacterial cell-volume of *P. gingivalis* wild-type but not Δ*gdhA* strain, an indicator of higher metabolism and growth. Taken together, this study highlights the pathophysiological importance of Glu for *P. gingivalis* growth-rate, biomass induction and survival in nutritionally limited host subcellular environments.

## Introduction

*Porphyromonas gingivalis* is a major Gram (-) opportunistic autochthonous bacterium of human oral mucosa. The oral mucosa houses a large variety of immune cells to provide first line of defense against pathogenic microorganisms where gingival epithelial cells (GECs) form the initial barrier and the important innate immune defense against colonizing microbes (1–3). *P. gingivalis* has adopted a variety of contextual mechanisms to evade host immune defenses and can effectively invade GECs to establish persistence in the gingival mucosa (4–10). *P. gingivalis* is a major pathogen in the etiology of advanced forms of periodontal diseases and more recently proposed to be involved in other chronic diseases progression (11–17). *P. gingivalis* has been shown to play key roles in oral microbial dysbiosis that results in selective enrichment of disease-causing microbes over beneficial microbes (18–22) in oral biofilm communities (23). On the other hand, *P. gingivalis’* effectual intraepithelial invasion via GECs provides an inner sanctum for the microorganism where *P. gingivalis* mainly survives and multiplies through the special formation of Endoplasmic Reticulum (ER)-rich autophagic vacuoles (7, 9, 24). The microorganism’s chronic intraepithelial colonization induces increased formation of microvasculature in the gingival epithelial mucosa that contributes to *P. gingivalis*’ systemic dissemination via the blood vessels and aids *P. gingivalis* with nutrients accessibility such as hemoglobin from ruptured vasculature due to periodontitis (25). The incapability of metabolizing saccharides renders *P. gingivalis* fastidious and heavily dependent on peptides/amino acids to obtain energy and micronutrients for growth (26–28). The genome sequence analysis further supports that *P. gingivali*s should have the capability of metabolizing several amino acids (27). However, only Serine as a single amino acid has been shown to be directly transported by *P. gingivalis* (29). Most of the reports showed that *P. gingivalis* consumes amino acids once they are liberated from short peptides and at least a dipeptide is required as a precursor to supply amino acids (30–34). However, all the previous studies were focused on either endpoint metabolites detection from planktonically grown *P. gingivalis* cultures to indirectly predict the type of amino acids *P. gingivalis* consumes. However, there has not been any detailed and direct growth kinetics studies to show the increase in growth of *P. gingivalis* when solely amino acids or peptides were used as nutrients. Moreover, the bacterial growth metabolism of intracellular *P. gingivalis*, especially in the double-membrane autophagic vacuoles where nutrients are limited, have been largely understudied (7, 35). In addition to providing metabolic energy and anabolic biomolecules for bacterial growth, some amino acids like Glutamate (Glu)/Glutamine (Gln) provide additional biological benefits. Because these amino acids, especially Glu is utilized as building block precursors for several important biomolecules like glutathione (GSH), a potent antioxidant (36). Interestingly, it has been shown that Gln is the key contributor of Glu in GSH synthesis (36) while we previously showed that intracellular *P. gingivalis* inhibits host antimicrobial reactive oxygen species (ROS) via increased GSH production by GECs (4, 8). Glu also serves as an essential precursor for the biosynthesis of a various of biologically important small molecules as well as nonessential amino acids (36, 37). Human cells can synthesize Glu through a variety of distinct metabolic pathways (38–43). However, Glu’s role as a critical nutrient exclusively for intracellular bacteria including *P. gingivalis* has not been known. *P. gingivalis* usurps host autophagic machineries to create a unique double membrane autophagic vacuoles for microbial replication and persistence in GECs (7, 24). Inhibition of autophagy by 3-methyladenine or depletion of ATG5 significantly reduces the viability of intracellular *P. gingivalis* (7) which indicates that *P. gingivalis* can efficiently scavenge nutrients from host to multiply and survive in the autophagic vacuoles (7, 24). However, most of the nutrients are in the forms of complex macromolecules like carbohydrates, lipids, and proteins, therefore, these nutriments are not easily accessible to the intracellularly invading pathogens. These microbes must have the machineries either to (i) degrade them to simpler molecules, (ii) utilize host metabolic machineries to do degrade them like by autophagy, or (iii) find simpler molecules already available in the niche. *P. gingivalis* produces proteases and dipeptidases that can degrade proteins or peptides to release amino acids that the microorganism can use for its metabolism (27, 44–46) in various extracellular host environments such as dental biofilm. However, the information on which amino acids *P. gingivalis* preferentially utilizes for intracellular host environments, specially, in the vacuolar life has been remained completely unexplored . Here we show for the first time that *P. gingivalis,* an opportunistic oral pathogen preferentially utilizes Glu as a key nutrient during the intracellular vacuolar life for replication and survival in GECs. We have specifically identified that *P. gingivalis* secretes Glutamate dehydrogenase (GdhA), a key enzyme encoded by PGN_1367 of *P. gingivalis*, for Glu metabolism determined by the selectively isolated intact *P. gingivalis*-containing vacuoles from the infected GECs (24). Treatment of GECs with Glu-Glu-dipeptide, the precursor of Glu, significantly increases the number *P. gingivalis* in the vacuoles and the size of the *P. gingivalis*-containing vacuoles significantly increases as well. To imitate nutritionally scarce environment as it is in these P*. gingivalis* replicative vacuoles, here we use a nutritionally reduced Trypticase-Soy-Broth (TSB) media having no additive hemin and confirm that Glu metabolism significantly supports and increases the growth of *P. gingivalis* in nutrient limited conditions. The importance of Glu metabolism for the bacterial vacuolar growth is further evident from our findings that a GdhA deletion mutant strain of *P. gingivalis* shows drastic growth defect both intracellularly as well as in typical microbiological *in vitro* culture systems. Since nutrient uptake is prerequisite to metabolism, we confirm that *P. gingivalis* uptakes Glu from the growth media. Interestingly, the super resolution Atomic-Force-Microscopy (AFM) analysis of *P. gingivalis* individual cells, further adds to the examination of *P. gingivalis*’ physiology that the cellular volume of *P. gingivalis* significantly increases in presence of Glu-Glu which indicates higher metabolic rates and that reconfirms higher metabolism of Glu is for *P. gingivalis* growth. These findings shed light on the biological relevance of Glu metabolism and *P. gingivalis* which could be a novel target for therapeutic intervention to control *P. gingivalis* infection associated chronic periodontitis and other chronic systemic diseases.

## Results

### Detection of glutamate catabolic enzyme glutamate dehydrogenase (GdhA) in the *P. gingivalis*-containing vacuoles from the infected human primary GECs

To explore which amino acids that *P. gingivalis* utilize during the bacterial intracellular survival and growth, we took a novel approach to sort out which *P. gingivalis* proteins were selectively enriched in *P. gingivalis*-containing autophagic vacuoles. Because this would help to map that protein(s) into metabolic pathways to understand its association with any amino acid or peptide metabolism. For this, we first performed unbiased label-free Liquid-Chromatography with Tandem-Mass-Spectrometry (LC-MS/MS) analysis of protein content in the intact *P. gingivalis*-containing vacuoles which were selectively isolated with a special method (as we recently described in (24). The label-free quantitative (LFQ) analysis of vacuolar proteins with over 95% confidence and intensity of spectra revealed that glutamate dehydrogenase (GdhA) enzyme encoded by PGN_1367 of *P. gingivalis* (Protein ID# B2RKJ1) were highly enriched in these vacuoles at all time points investigated and highest intensity of GdhA was found at 6 h with an LFQ value of 26 ± 3 (**Fig. 1A**). To further validate this LC-MS/MS result, we next tested the presence of GdhA by classical immunofluorescence (IF) staining. The isolated vacuoles were stained with the *P. gingivalis* recombinant GdhA, rGdhA (PGN_1367) -targeted anti-rabbit antibody prepared in this study and the results showed that GdhA was specifically present at high intensity (red fluorescence). The presence of *P. gingivalis* was also visualized using *P. gingivalis*-specific anti-mouse antibody in the same sample (green fluorescence). The staining of the vacuole by thiol-tracker (blue fluorescence) was used to confirm the intactness of the isolated vacuoles with *P. gingivalis* (**Fig. 1B**).

**Fig. 1.**
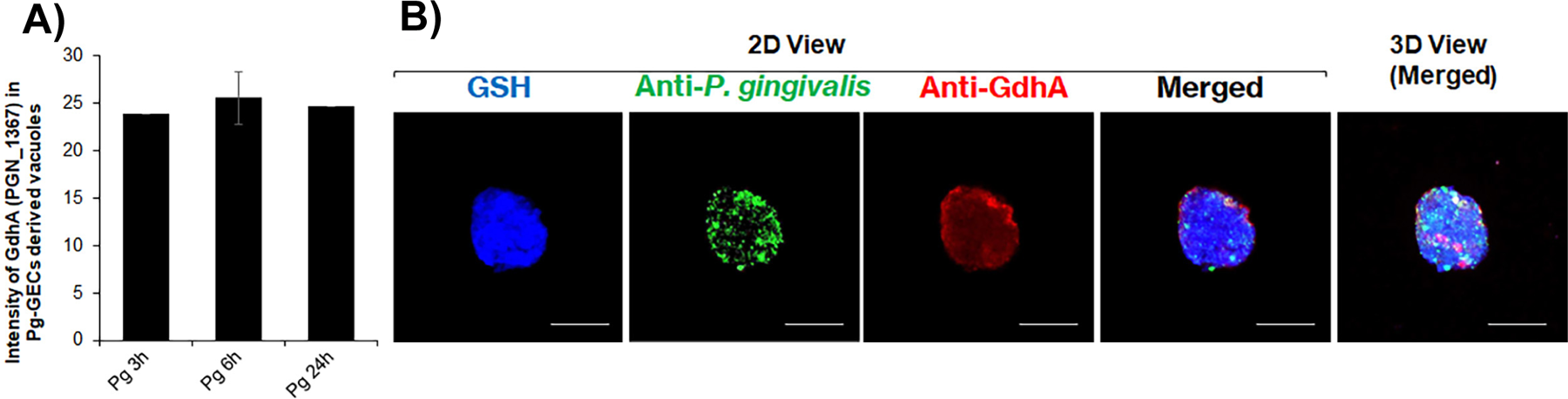
High peak intensity of glutamate dehydrogenase (GdhA) of *P. gingivalis* in autophagic vacuoles derived from the infected GECs. Following infection of GECs with wild-type *P. gingivalis*, intact autophagic double membrane vacuoles containing the bacteria were isolated using the special protocol (as we recently described in. (24)). The vacuolar proteins were then analyzed by a high resolution and high mass accuracy LC-MS/MS instrument. The label-free quantities (LFQ) of GdhA (PGN_1367, protein ID# B2RKJ1)-specific peptides were detected with over 95% confidence based on spectrum intensity and Blast search. The assay was performed twice with at least 6-replicates pooled together in each assay and the data is represented as Mean ± SD (A). For immunofluorescence (IF) staining of vacuoles, GECs were treated with 2 mM Glu-Glu exogenously for 1 h prior to infection with wild-type *P. gingivalis*. Vacuoles were isolated, fixed and stained with *P. gingivalis*-specific antibody and GdhA-specific antibodies to detect *P. gingivalis* (green) and GdhA (red), respectively, and integrity of the vacuoles were confirmed by staining with thiol-tracker as blue. The images were captured by a confocal microscope (Leica DM6 CS Stellaris 5 Confocal/Multiphoton System) at 63x magnification where the scale bar is 5 µm. The images were then processed and analyzed by Imaris software. Representative images of vacuoles of three independent assays are shown (B).

### Glutamate is preferred than glutamine by the vacuolar *P. gingivalis* in human primary GECs and molecular docking showed GdhA putatively used glutamate as substrate

Since host cells consume glutamine (Gln) for growth and produces Glu (36), we investigated whether *P. gingivalis* infection had any impact on the GECs’ Gln to Glu conversion and whether *P. gingivalis* prefers Gln over Glu in GECs. We performed the Gln to Glu conversion assay at first with GECs infected by *P. gingi*valis over 24 h and interestingly, it showed that *P. gingivalis* significantly (P<0.01) induced the conversion of Gln to Glu and this conversion level was highest at 6 h of infection and addition of exogenous Gln further increased the Gln to Glu conversion level (**Fig. 2A**). We then tested this conversion analysis specifically with the isolated *P. gingivalis*-containing vacuoles derived from GECs-infected for 6 h with *P. gingivalis* and as anticipated, Gln to Glu conversion was significantly (P<0.01) induced (**Fig. 2B**). These findings indicate that *P. gingivalis* prefers Glu and therefore induces Gln to Glu conversion to make more Glu available to potentially be utilized for the bacterial growth. Since GdhA was highly present in the *P. gingivalis*-containing vacuoles and Glu is the substrate of GdhA, we performed molecular docking to predict the potential binding of Glu with GdhA. The docking assay showed that Glu (sphere shape) fits into the substrate binding pocket consisting of K88, K112, and S378 (pink color) of GdhA (**Fig. 2C**). A GdhA homolog Glud produced by *C. difficile* is well studied since Glud is highly secreted and hence it is used as an indicator of *C. difficile* infection in stool samples (47, 48), we superimposed 3D structures of GdhA with Glud and the result showed high similarities in 3D structures with an RMSD (root mean square deviation) value of 1.103[ (**Fig. 2D**). The alignment of amino acid sequences of these two proteins revealed that the Glu binding sites (K88, K112, and S378 pink color in GdhA), active sites (K124 and D164 in GdhA, red color), and cofactor/coenzyme NAD^+^ binding sites (T207 and N238 in GdhA, blue color) were highly conserved (**SI Fig. 1**). Based on our findings and KEGG pathway analysis, we schematically mapped the GdhA on Glu catabolic pathway that shunts the intermediates to TCA cycle and makes carbon and nitrogen available which are necessary for the growth of *P. gingivalis* (**Fig. 2E**).

**Fig. 2.**
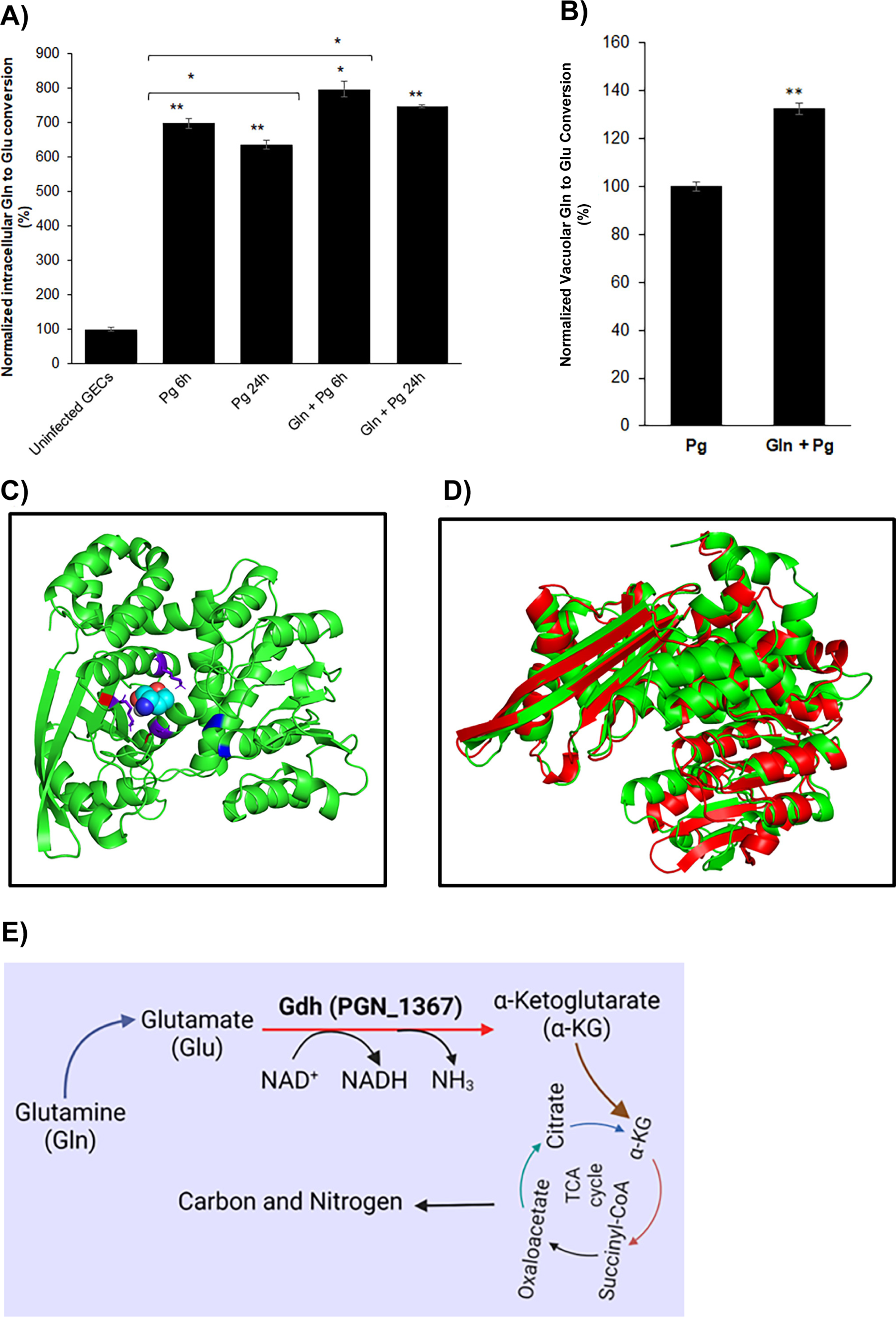
Induction of intra/sub-cellular glutamine (Gln) to glutamate (Glu) conversion by *P. gingivalis* in GECs and molecular docking of glutamate catabolic enzyme Glutamate dehydrogenase. GECs were infected with wild-type *P. gingivalis* (Pg) for 6 h or 24 h with or without 2 µM exogenous Gln. GECs or the vacuoles were isolated to measure (A) intracellular or (B) intra-subcellular (i.e., vacuolar) conversion of Gln to Glu using Gln/Glu-Glo assay kit. The data is presented as Mean ± SD and Students’ two-tailed T-TEST was used for statistical analysis where a p-value of <0.01 (**) was considered statistically significant. The assay was performed twice with at least 6-replicates in each assay. (C) Molecular docking between GdhA (Cartoon shape) and its substrate Glu (Sphere shape) was analyzed by supercomputer at ClusPro which was then analyzed by PyMol software. The Glu binding sites in GdhA are shown in purple color, active sites in Red color, and its cofactor NAD+ binding sites in Blue color. (D) GdhA (Green) and GluD (Red) are glutamate dehydrogenase in *P. gingivalis* and *Clostridium difficile*, respectively. They were superimposed using PyMol molecular docking software. (E) GdhA catabolic reaction showing Glu as a potential source of carbon and nitrogen for *P. gingivalis* growth.

### GdhA is secreted from *P. gingivalis* and possesses canonical Gdh activity

The GdhA secretion from *P. gingivalis* were validated by both western blot and ELISA using the *P. gingivalis* rGdhA-specific antibody prepared in this study. The bacterial cells free supernatant (CFS) was prepared from *P. gingivalis* wild-type (WT) or Δ*gdhA* strains following standard protocol. A 10 ng of purified rGdhA (expressed and purified in this study as detailed in the methods section) was used as a positive control. The immunoblot analysis showed that the GdhA was abundantly present in the CFS of WT strains but not at all of the Δ*gdhA* strain. Interestingly, the level of GdhA was significantly increased (more than 2.5-fold) in the presence of Glu-Glu-dipeptide (**Figs. 3A and 3B**). Since ELISA is more sensitive than immunoblot and can estimate the exact amount of protein of interest present in the sample, we employed ELISA with Rabbit anti-rGdhA specific primary antibody coupled with HRP-conjugated mouse secondary monoclonal antibody (only specific for the Fc of rabbit IgG) to reconfirm the secretion of GdhA from *P. gingivalis* and to accurately quantify the level of GdhA in the CFS. The results displayed, the absorbance for CFS of Δ*gdhA* strain was almost the same as blank sample and hence this value was subtracted from the values of WT strains. The amount of GdhA was estimated as 644 ± 15 ng/mL in the CFS of WT strain, and the amount was increased by 2.3 ± 0.3-fold to 1456 ± 167 ng/mL in presence of Glu-Glu-dipeptide (**Fig. 3C**). The ELISA data were in line with the immunoblot result in terms of fold increase with Glu-Glu supplementation. Like blank sample and CFS of Δ*gdhA* strain, no reactivity of rGdhA antibody was observed even with very high concentration (5 µg/mL) of BSA. These results demonstrated that GdhA was notably secreted from the *P. gingivalis* WT strain and Glu-Glu-dipeptide supplementation significantly induced the level of GdhA production from *P. gingivalis*. Both immunoblot and ELISA data corroborated the LC-MS/MS analysis of GdhA presence in the *P. gingivalis*-containing vacuoles derived from the infected GECs. The absence of any bacterial cells in the CFS was confirmed by observing no visible growth when CFS was inoculated TSB or TSB blood agar plate (Data not shown). Moreover, *P. gingivalis*’ 16S rRNA-specific PCR was negative to detect any DNA of *P. gingivalis* (Data not shown). Alternatively, this underscores that the release of GdhA from lysed cells into the CFS was very unlikely.

**Fig. 3.**
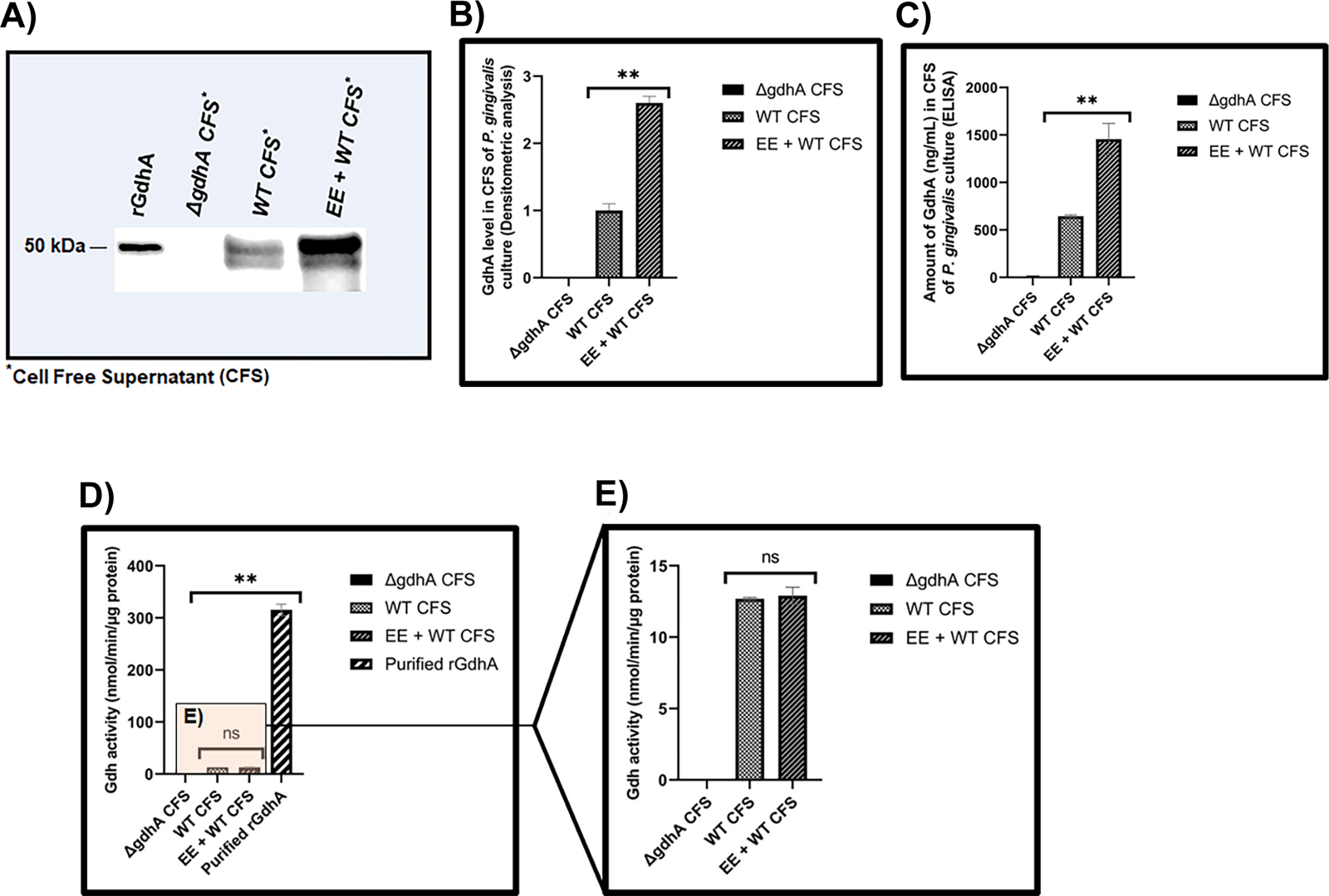
GdhA (PGN_1367) of *P. gingivalis* indeed is a secretory protein and possesses typical GdhA activity. A 200 mL culture of *P. gingivalis* wild-type (WT) strain with/without 2 mM Glu-Glu-dipeptide and a 400 mL culture of isogenic *gdhA* deletion mutant (Δ*gdhA)* strain were grown to mid exponential phase (OD_600_ ≤ 0.8). Bacterial cells were removed by centrifugation. The supernatant was filtered (0.1 µm pore size) to make it cell free supernatant (CFS) and proteins were precipitated by salting-out (about 80% saturation) followed by centrifugation. The precipitate was resuspended in 5 mL of cell lysis buffer. The samples were dialyzed against 400x excess PBS buffer at 4°C overnight with two changes of the PBS buffer. The samples were then concentrated using Amicon 10-kDa cutoff filter system to 1 mL. Protein concentration was estimated by Bradford assay. A 40 µg of protein of CFS samples and 10 ng of purified rGdhA were used for western blotting with anti-rabbit rGdhA antibody (A). Image J software was used for densitometric analysis of the western blots for GdhA (B). The amount of GdhA in the CFS samples were also estimated by ELISA using anti-rabbit rGdhA antibody and Mouse anti-rabbit IgG Fc monoclonal antibody conjugated with HRP. Purified rGdhA was used to prepare the standard curve which was used to calculate the amount of GdhA in the CFS samples (C). The CFS samples (about 50 µg total protein present in the CFS) as well as purified rGdhA (1.0 µg) were also used to test GdhA activity using glutamate as substrate by modified Gln/Glu-Glo assay kit. The activity was expressed as amount of glutamate (nanomole) catalyzed by 1.0 µg of protein by 1.0 min (D). The Gdh activity graph with only CFS samples were zoomed in (E). The data is presented as Mean ± SD and Students’ two-tailed T-TEST was used for statistical analysis where a p-value of <0.01 (**) was considered statistically significant. The assay was performed twice with at least 3 replicates in each assay. Here, “ns” indicates “not significant”.

We next investigated the biochemical activity of the GdhA secreted from *P. gingivalis* in the CFS as well as the purified rGdhA. Since Gdh utilizes NAD^+^ as a coenzyme and catalyzes the conversion of glutamate to α-ketoglutarate and NADH, we adapted the glutamine/glutamate-Glo kit to measure the activity of Gdh in terms of how much of the glutamate is consumed overtime. The activity assay revealed that CFS of both *P. gingivalis* WT strain with and without Glu-Glu had similar activity 13 ± 0 and 13 ± 1 nmol/min/μg of total protein present in the CFS and no Gdh activity was found in the CFS of the Δ*gdhA* strain (**Figs. 3D and 3E**). The purified rGdhA used as a control had significantly (∼25-fold) more activity of 315 ± 11 nmol/min/μg protein than the CFS (**Fig. 3D**) and the kit provided Gdh showed 10-times higher activity of 3057 ± 98 nmol/min than the rGdhA of *P. gingivalis* (data not shown). Since the Gdh provided with the assay kit was proprietary of the company (Promega) and the amount of protein and its source were not provided, we could not exactly compare its activity with the purified rGdhA prepared in this study.

### Glutamate catabolism increased intracellular level of *P. gingivalis* wild-type strain but not GdhA deficient isogenic mutant strain, Δ*gdhA* in GECs

Published studies indicated that *P. gingivalis* needs at least dipeptide for the bacterial growth since single amino acid does not significantly impact on its growth (28, 30, 31, 33, 49, 50). We also tested different concentrations (10 mM, 20 mM, and 40 mM) of free Glu in bacterial culture media and reconfirmed that it did not increase the growth of *P. gingivalis* (**Figs. 8 and 9**). For this reason, we used Glu-Glu-dipeptide as Glu precursor. To decipher the direct roles of Glu metabolism in the intracellular growth of *P. gingivalis*, GECs were treated with Glu-Glu prior to the infection with *P. gingivalis* wild-type or Δ*gdhA* strains. The intracellular level of metabolically active *P. gingivalis* was then quantified by classical antibiotics protection assay coupled with 16S rRNA-based qPCR. The results clearly showed that Glu-Glu treatment significantly increased the level of intracellular *P. gingivalis* wild-type strain over the time and being highest at 24 h (**Fig. 4A**). In contrast, level of Δ*gdhA* strain was significantly (P<0.01) decreased upon Glu-Glu treatment (**Fig. 4B**). The overall bacterial population was about 2-log less in case of Δ*gdhA* strain compared to the wild-type strain (**Fig. 4A-B**).

**Fig. 4.**
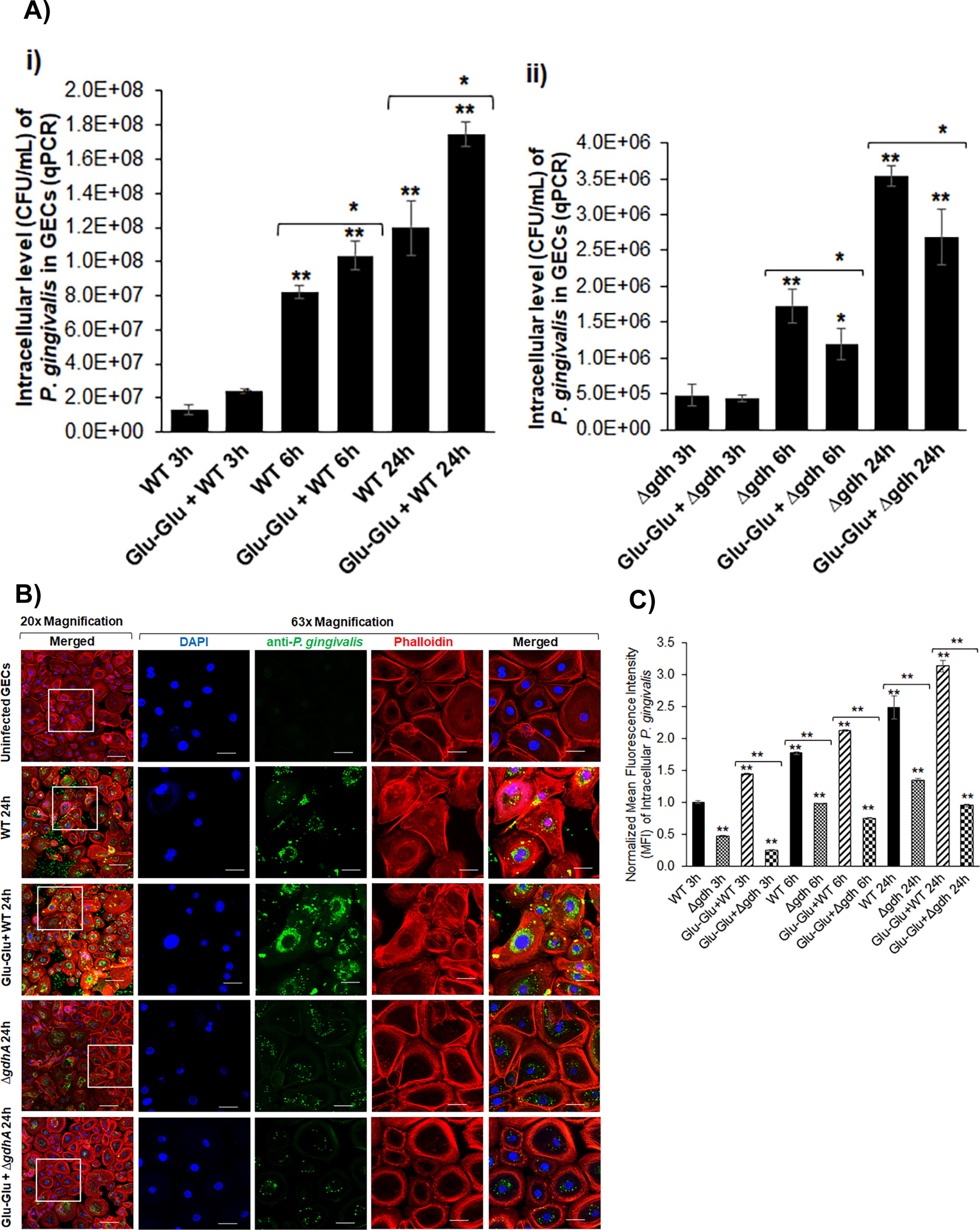
Intracellular level of *P. gingivalis* wild-type (WT) and Δ*gdhA* strains in GECs. Following infection of GECs grown in 6-well plates (A) or in 4-well chamber slides (B) with WT or Δ*gdhA* strains of *P. gingivalis* (Pg) in presence or absence of 2 mM Glu-Glu for 3 h, 6 h, and 24h, samples were treated with antibiotics to kill extracellular bacteria (A) or fixed and subjected to IF staining (B). For (A) GECs with WT (i) or Δ*gdhA* (ii) strain infections were collected with Qiazol and RNA was isolated. cDNA was synthesized from isolated RNA and Pg was quantified by qPCR using 16S RNA primers specific to Pg. For (B), fixed cells were stained with anti-rabbit *P. gingivalis-*specific antibody and a goat anti-rabbit Alexa fluor 488-conjugated secondary antibody to stain Pg as green. Rhodamine Phalloidin was used to stain the cells cytoskeleton as red. The samples were washed and mounted with coverslips using antifade mounting medium with DAPI (to stain nuclei as blue) and visualized under a confocal microscope (Leica DM6 CS Stellaris 5 Confocal/Multiphoton System) at 20x (100 µm scale bar)/63x (20 µm scale bar) magnification. Images representative of 24h infection of at least two separate experiments performed in duplicates are shown here (B). The mean intensity of green fluorescence of at least 25 cells of each indicated condition derived from intracellular Pg by IF staining (B) was quantified (C). The data in (A) or (C is represented as Mean ± SD of two/three replicates each assayed in duplicates/triplicates. Students’ two-tailed T-TEST was used for statistical analysis and a p-value of <0.05 (*) or <0.01 (**) was considered as statistically significant.

To further validate the antibiotic protection assay data, we performed IF staining of intracellular *P. gingivalis* following infection of GECs with wild-type or Δ*gdhA* strains over 24 h. Visual observation of IF images showed higher level of intracellular wild-type strain in GECs with Glu-Glu treatment compared to without Glu-Glu in wild-type *P. gingivalis* (**Fig. 4C**). In contrast, as observed in antibiotic protection assay, intracellular level of Δ*gdhA* was very low compared to wild-type strain and Glu-Glu treatment further decreased Δ*gdhA* level (**Fig. 4C**). Measurement of mean fluorescence intensity (MFI) of green fluorescence (*P. gingivalis*) corroborated the visual observations of IF images. For example, Glu-Glu treatment increased the MFI of wild-type strain from 2.5 ± 0.2 to 3.1 ± 0.1 (p<0.1), whereas it decreased from 1.3 ± 0.0 to 0.9 ± 0.0 in Δ*gdhA* strain (**Fig. 4D**). Collectively, these results suggested that Glu plays key roles in intracellular replication of *P. gingivalis* and GdhA is vital for *P. gingivalis* to catabolize Glu.

### Glu played significant roles via GdhA in size and integrity of *P. gingivalis*-containing vacuoles in GECs

Since *P. gingivalis* rapidly traffics into double-membrane autophagic vacuoles (7) for replication and survival in GECs and GdhA was detected in the vacuoles as shown above, we then examined the effect of Glu on the integrity and size of isolated *P. gingivalis*-containing vacuoles for *P. gingivalis* wild-type or Δ*gdhA* strains following staining for thiol (e.g., glutathione, GSH). Interestingly, Glu-Glu treatment drastically increased the size of the *P. gingivalis* wild-type strain-containing vacuoles as it is depicted from both 2D, 3D, and orthogonal confocal microscopy views of vacuoles (**Fig. 5A-B**) and the vacuoles were fully intact. Glu-Glu treatment with *P. gingivalis* wild-type infection increased the width and length of the vacuoles by 2.2 ± 0.3 and 2.3 ± 0.4-fold, respectively, (**Fig. 5C**). Additionally, the visible level of *P. gingivalis* wild-type in the vacuoles were higher in Glu-Glu treated samples. In contrast, the vacuoles were either not finely formed or disintegrated in the isogenic Δ*gdhA* strain infected samples and the bacteria-containing vacuoles were severely disintegrated upon Glu-Glu treatment as it is shown in the orthogonal view (**Fig. 5B**). The size (width or length) of the vacuoles with *P. gingivalis* Δ*gdhA* strain infection were more than 2-fold smaller compared to wild-type strain. The Glu-Glu treatment further decreased the size of the Δ*gdhA* strain infected vacuoles by 2-fold (**Fig. 5C**).

**Fig. 5.**
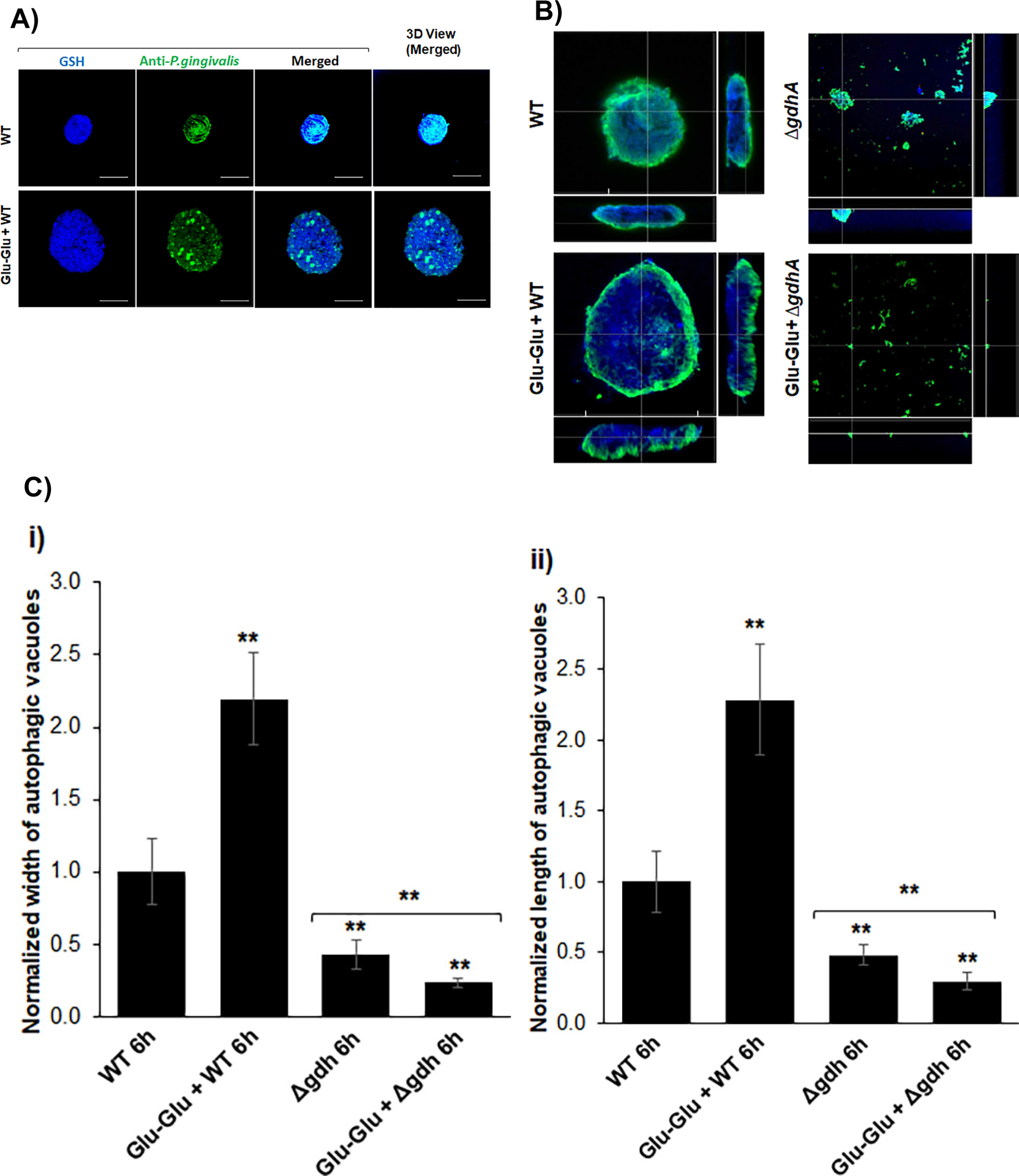
Size of *P. gingivalis*-containing vacuoles derived from *P. gingivalis*-infected GECs upon exogenous Glu treatment. GECs were pre-treated with 2 mM Glu-Glu for 1 h prior to infection with wild-type (WT) and Δ*gdhA* strains of *P. gingivalis* for 6h. Vacuoles were isolated, fixed, and stained with *P. gingivalis*-specific antibody to detect *P. gingivalis* (green) and stained with thiol-tracker to stain the thiol/glutathione (GSH) of vacuoles. Images were captured by a confocal microscope (Leica DM6 CS Stellaris 5 Confocal/Multiphoton System) at 63x magnification where the scale bar is 5 µm. The images were then processed and analyzed by Imaris software. (A) 2D and 3D views of representative images of vacuoles derived from GECs infected with WT Pg. (B) Zoomed Orthogonal view of representative images of WT Pg or Δ*gdhA* Pg-infected vacuoles. (C) The size (width [i] and length [ii]) of at least 25 vacuoles from three independent assays was quantified and normalized to WT 6 h condition. The data is represented as Mean ± SD and Students’ two-tailed T-TEST was used for statistical analysis where a p-value of <0.01 (**) was considered as statistically significant.

### *P. gingivalis* taken-up Glu which is pre-requisite to catabolism and preferentially utilized Glu-Glu-dipeptide in bacterial culture media

Since Glu-Glu-dipeptide, significantly increased the size of live *P. gingivalis*-containing vacuoles along with the intracellular bacterial growth in GECs, we hypothesized that *P. gingivalis* might uptake Glu from the environment. To investigate that we used radiolabeled L-[^14^C(U)] Glu. Because use of radiolabeled Glu will help to understand whether it has been taken-up by the bacterial cells or not, though supplementation of free Glu did not show any increase in *P. gingivalis* growth. Similar study was done before but with radiolabeled Serine (29). Our results illustrated that *P. gingivalis* can successfully take-up ^14^C-Glu when grown in TSB or TSBNH media where both media were supplemented with ^14^C-Glu and the uptake amount was observed to increase over the time. Moreover, the ^14^C-Glu uptake *by P. gingivalis* was significantly higher in TSBNH media compared to TSB, underlining the potential significance of Glu in nutritionally reduced conditions (**Fig. 6A**). Next, we investigated if other dipeptides that can supply amino acids other than Glu could provide any similar or better growth than Glu-Glu. Since ^14^C-Glu uptake was more in TSBNH media, this reduced media was used to test growth kinetics of *P. gingivalis* in different dipeptides. Among four individual dipeptides tested in TSBNH media, the Glu-Glu supplemented media provided the best growth to *P. gingivalis*. The order of dipeptides based on growth kinetics of *P. gingivalis* was Glu-Glu>Glu-Gln>Gln-Gln>Arg-Arg (**Fig. 6B**). This finding further supports why Glu catabolic enzyme, GdhA was highly enriched and Glu was made available from Gln instead of directly catabolizing Gln by *P. gingivalis* in the autophagic vacuoles.

**Fig. 6.**
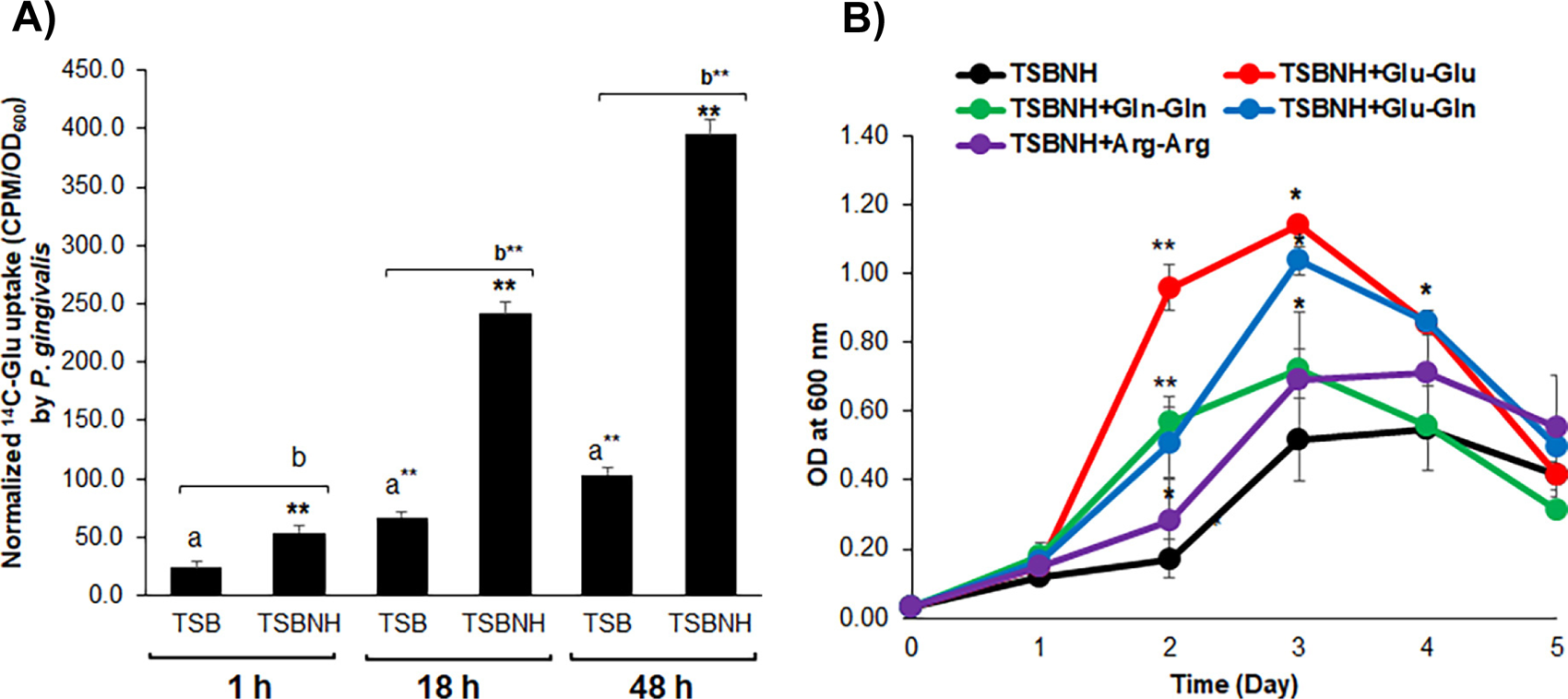
Glutamate uptake by wild-type *P. gingivalis* and growth kinetics upon supplementation with different dipeptides including Glu-Glu. *P. gingivalis* wild-type strain was incubated in TSB and reduced TSB media with no added hemin (i.e., TSBNH) with or without radiolabeled L-[^14^C(U)] Glu (1.0 mCi/mmol) at 37°C under anaerobic conditions. Bacterial growth was monitored by measuring OD at 600 nm and 1 mL of the culture was collected at 1 h, 18 h, and 48 h time points to harvest bacterial cells by centrifugation at 10,000 x *g* for 5 min. Bacterial cells pellet was washed three times with 1 mL PBS to remove any ^14^C-Glu outside of the cells. Cells were then lysed with 20 mg/ mL of lysozyme and mixed with liquid scintillation cocktail before measuring the radioactivity (Counts per minute [CPM]) using a liquid scintillation counter. Radioactivity reading of bacterial culture without ^14^C-Glu was used as blank and the value was subtracted from the corresponding readings of bacterial culture with ^14^C-Glu. Glutamate uptake by *P. gingivalis* was normalized to OD_600_ of *P. gingivalis* growth and, hence, expressed as CPM/OD_600_. The data is represented as Mean ± SD, where n = 3; and a p-value of <0.01 was considered statistically significant (Students’ two-tailed T-Test). Here, a and b indicate T-Test among 1 h,18 h, and 48 h of TSB and TSBNH, respectively. Results of an assay out of three, each assayed in triplicates are shown here (A). For growth analysis (B), *P. gingivalis* wild-type strain was grown in TSBNH supplemented with 10 mM dipeptide of Glu-Glu, Glu-Gln, Gln-Gln, or Arg-Arg and the growth was monitored everyday over a 5-day period by measuring the optical density (OD) of culture turbidity at 600 nm. The data is represented as Mean ± SD of three replicates of an assay out of three assays. Students’ two-tailed T-TEST was used for statistical analysis and a p-value of <0.05 (*) or <0.01 (**) was considered statistically significant (B).

### Glu-Glu dipeptide supplementation increased *P. gingivalis* growth in bacterial culture media especially in TSB reduced media

Since Glu-Glu-dipeptide provided highest growth of *P. gingivalis* in bacterial broth culture media among other amino acids as well as Glu-Glu substantially enhanced the intracellular vacuolar growth of *P. gingivalis* in GECs, we wanted to further resolve the intracellular growth trajectories in the reductionist bacterial culture media models with biological relevance. At first growth of *P. gingivalis* wild-type strain in regular TSB (hemin 5 µg/mL) with or without Glu-Glu was investigated. Stimulation by Glu-Glu significantly increased the growth of *P. gingivalis* and it was about 1.3-fold higher (OD_600_: 2.5 ± 0.0 Vs to 1.9 ± 0.1) in TSB + Glu-Glu than TSB alone respectively (**Fig. 7A**). Since TSB is a rich media and *P. gingivalis* reaches its maximum growth peak much rapidly, use of nutritionally limited TSB media, like TSBNH that slows down the bacterial growth might resolve better contribution of any other supplements when added with TSBNH. Glu-Glu supplementation in TSBNH significantly contributed to the increased growth of *P. gingivalis*. At day-3, the growth turbidity at OD_600_ was 3.2-fold higher in presence of Glu-Glu than without Glu-Glu in TSBNH (0.89 ± 0.04 Vs 0.28 ± 0.06) (**Fig. 7B**). Since the fold change of *P. gingivalis’* growth in TSBNH media was higher than TSB when supplemented with Glu-Glu, TSBNH media was used for other experiments unless otherwise specified. The Δ*gdhA* strain of *P. gingivalis* showed significant growth defect even when grown in nutrient rich TSB media and the growth at maximum peak at day-2 was 0.45 ± 0.05 compared to 2.5 ± 0.0 of wild-type strain (**Fig. 7C**). Supplementation of Glu-Glu further significantly (p<0.01) decreased the growth of the Δ*gdhA* strain from 0.45 ± 0.05 to 0.15 ± 0.03. The Δ*gdhA* strain failed to grow in TSBNH media (**Fig. 7C**). Moreover, the lag time was significantly (48 hours) higher compared to wild-type strain (Data not shown). It is to clarify that in the growth assay (as detailed in methods and materials section), both wild-type and Δ*gdhA* strains were pre-grown and concentrated to have about same start-up OD (about 0.03) in the beginning of each growth assay. Therefore, it is not possible to determine the lag time from the growth curves presented here.

**Fig. 7.**
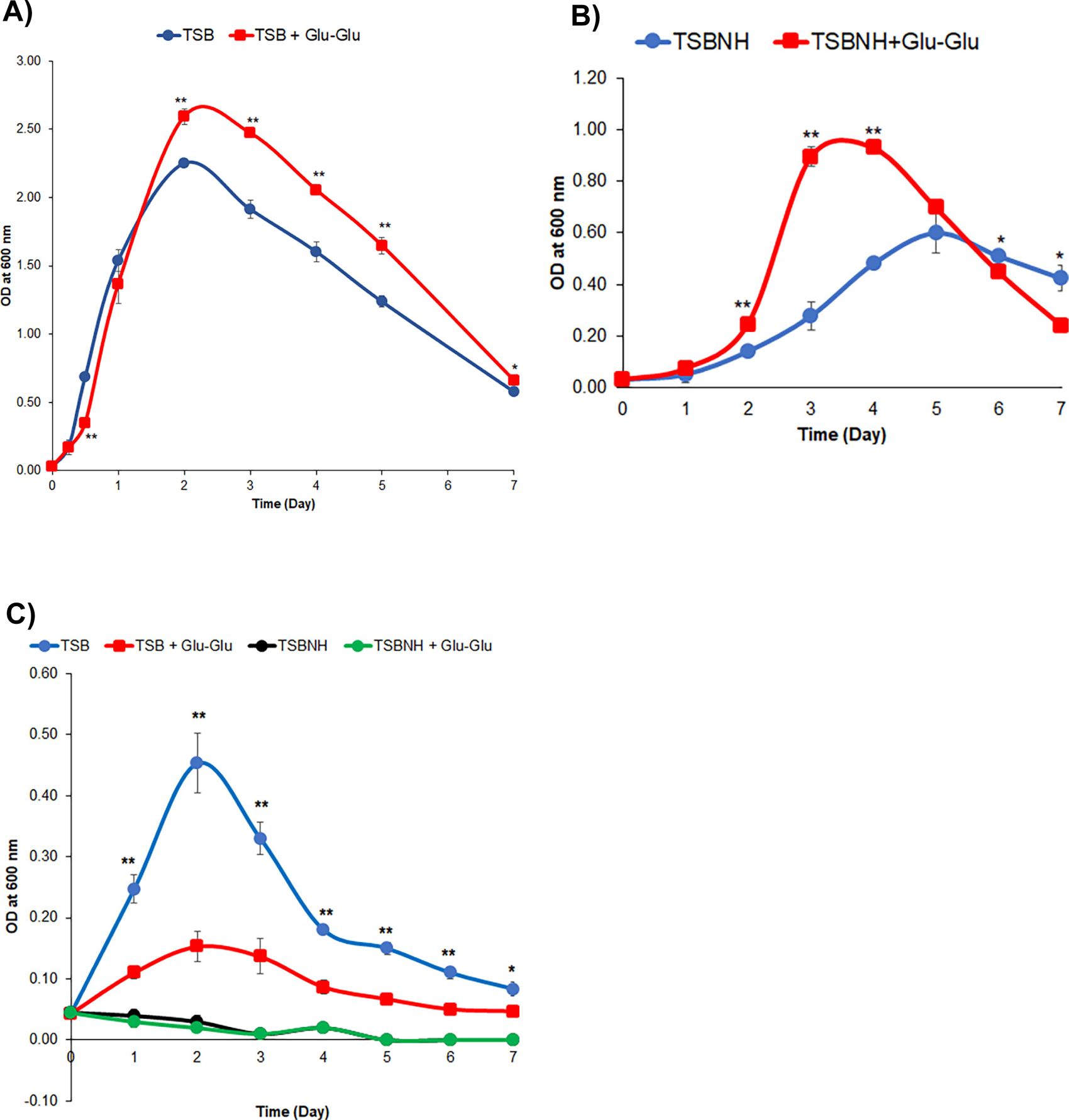
Growth of *P. gingivalis* wild-type and Δ*gdhA* strain*s* in regular TSB or reduced TSB (TSBNH) media supplemented with Glu-Glu-dipeptide. *P. gingivalis* wild-type (WT) or Δ*gdhA* strains were grown in either TSB or TSBNH media supplemented with 10 mM Glu-Glu-dipeptide. The growth was monitored at indicated time points over a 7-day period by measuring optical density (OD) of culture turbidity at 600 nm. Growth of WT in TSB (A) or TSBNH (B) were separately plotted, whereas growth of Δ*gdhA* in TSB or TSBNH were plotted together (C) to reveal that Δ*gdhA* strain did not grow in TSBNH media. The assays were repeated at least three times each assayed in triplicates and the results of a representative assay are shown here. The data is represented as Mean ± SD of three replicates. Students’ two-tailed T-TEST was used for statistical analysis and a p-value of <0.05 (*) or <0.01 (**) was considered statistically significant.

### Glu supplementation only after priming the growth with Glu-Glu dipeptide synergistically increased the growth of *P. gingivalis*

As described above, free Glu supplementation did not show any growth increase in *P. gingivalis*. Then we thought adding free “Glu” after priming the growth of *P. gingivalis* with Glu-Glu dipeptide first might enhance the bacterial growth. Therefore, to investigate the synergistic effect of Glu-Glu and free “Glu” in a stepwise manner, a similar growth assay was first performed in TSB media. Following 6 h (0.25 Day) of post-inoculation growth in the media with/without Glu-Glu dipeptide, 40 mM amino acid “Glu” was added. The results showed that “Glu-Glu +Glu” significantly (P<0.1) increased the growth of *P. gingivalis* wild-type strain especially at Day-3 compared to Glu-Glu alone as revealed by both OD measurement as well as by colony counts (**Fig. 8**). The isogenic *P. gingivalis* Δ*gdhA* strain did not show any growth increase with “Glu-Glu + Glu” supplementation, instead it showed more growth decrease compared to Glu-Glu alone (Data not shown). The assay was then repeated in TSBNH media for wild-type strain only (as Δ*gdhA* strain failed to grow in TSBNH media). After overnight growth in TSBNH with/without Glu-Glu dipeptide, 40 mM “Glu” was added. As predicted, “Glu” supplementation after priming the growth with Glu-Glu dipeptide showed significantly higher growth turbidity at OD 600 nm than Glu-Glu supplementation alone (1.2 ± 0.0 Vs 0.9 ± 0.0 at Day-3) (P<0.01, **Fig. 9A**). The representative picture of culture tubes was also captured at Day-3, the time point at which *P. gingivalis* reached its highest growth under the condition investigated. The image shows that the growth differences of *P. gingivalis* were highly visible in the culture tubes and the wild strain bacteria grown in presence of Glu-Glu dipeptide or Glu-Glu + Glu were markedly more turbid compared to TSBNH or TSBNH + Glu (**Fig. 9B**). To quantify metabolically active *P. gingivalis* in the same cultures, 16S rRNA-based qPCR was carried out with Day-2 to Day-4 cultures. The qPCR assay revealed the presence of higher live number of *P. gingivalis* when grown in presence of Glu-Glu + Glu than in Glu-Glu alone TSB media, and as it was shown, Glu-Glu treatment itself showed higher number of bacterial cells compared to TSBNH or TSBNH + Glu respectively (**Data not shown**). Next, we plated the serially diluted cultures on TSB blood agar to count live and culturable bacterial numbers. As hypothesized, supplementation of both Glu-Glu dipeptide and free “Glu” showed significantly (P<0.01) higher number of bacterial cells compared to Glu-Glu alone at Day-2 to Day-4, being highest at Day-3 (5.1E+09 ± 6.0E+08 Vs 3.6E+09 ± 5.3E+08). Similarly, Glu-Glu itself also showed higher number (P<0.01) of bacterial cells when compared to TSBNH alone at Day-3 and -4 (**Fig. 9D**). Taken together, the result of live bacterial count assay further supports that the increase in growth turbidity measured by OD_600_ was due to the presence of a higher number of viable *P. gingivalis* cells.

**Fig. 8.**
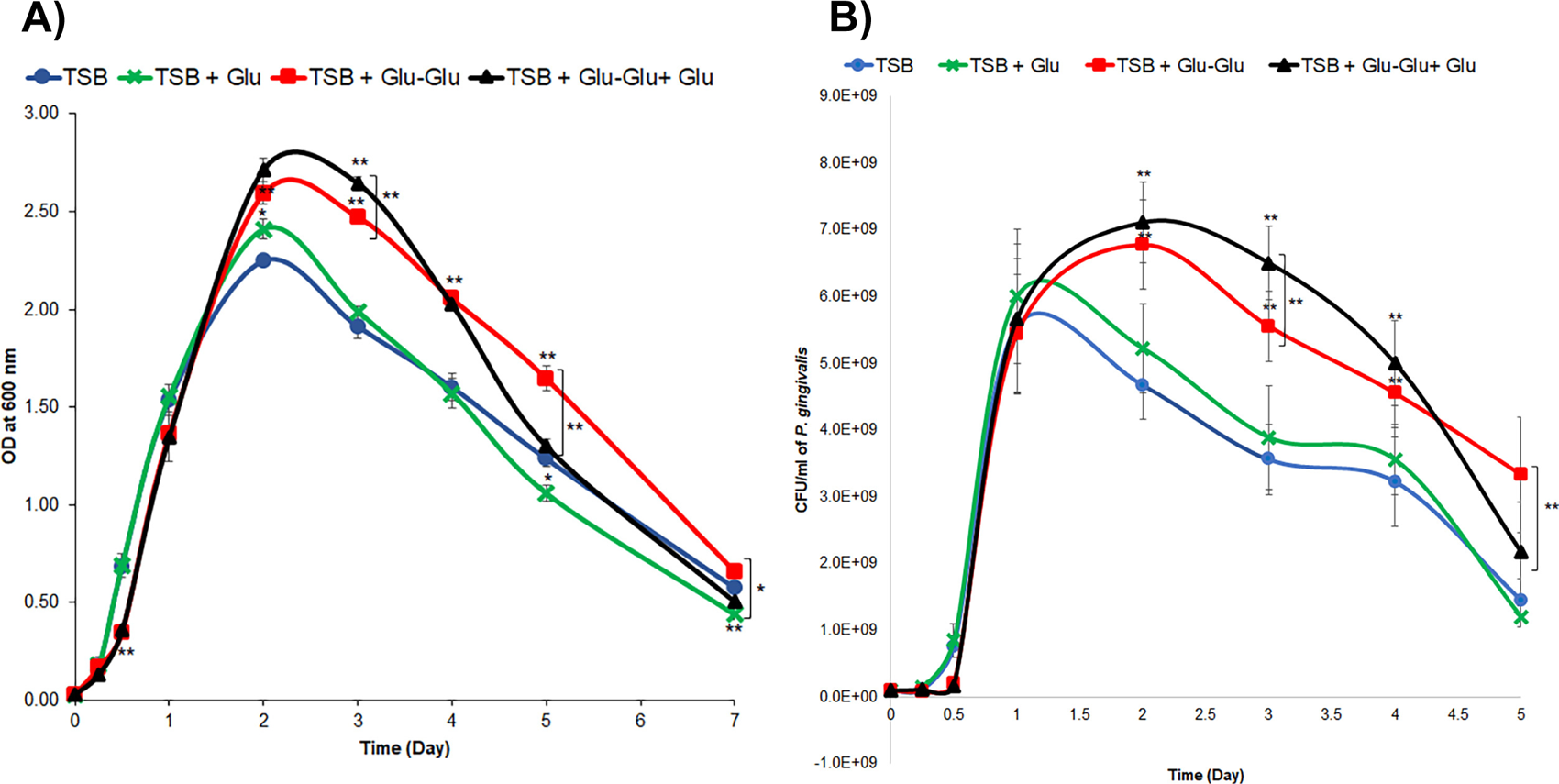
Growth of *P. gingivalis* wild-type strain in regular TSB media supplemented with Glu-Glu-dipeptide and free amino acid glutamate. *P. gingivalis* wild-type (WT) strain was grown in TSB media supplemented with 10 mM Glu-Glu-dipeptide followed by addition of 40 mM glutamate at Day-0.25. The growth was monitored every 0.25-day (6 h) or 1 day (24 h) interval whenever appropriate over a 7-day period by measuring optical density (OD) of culture turbidity at 600 nm (A). The growth of WT strain was also monitored over 5 days by CFU count by drop (10 µl x 3) plating of 10-fold series diluted samples on TSB blood agar (B). The data is represented as Mean ± SD of three replicates. Students’ two-tailed T-TEST was used for statistical analysis and a p-value of <0.05 (*) or <0.01 (**) was considered statistically significant. The assays were repeated at least three times each assayed in triplicates and the results of a representative assay are shown here.

**Fig. 9.**
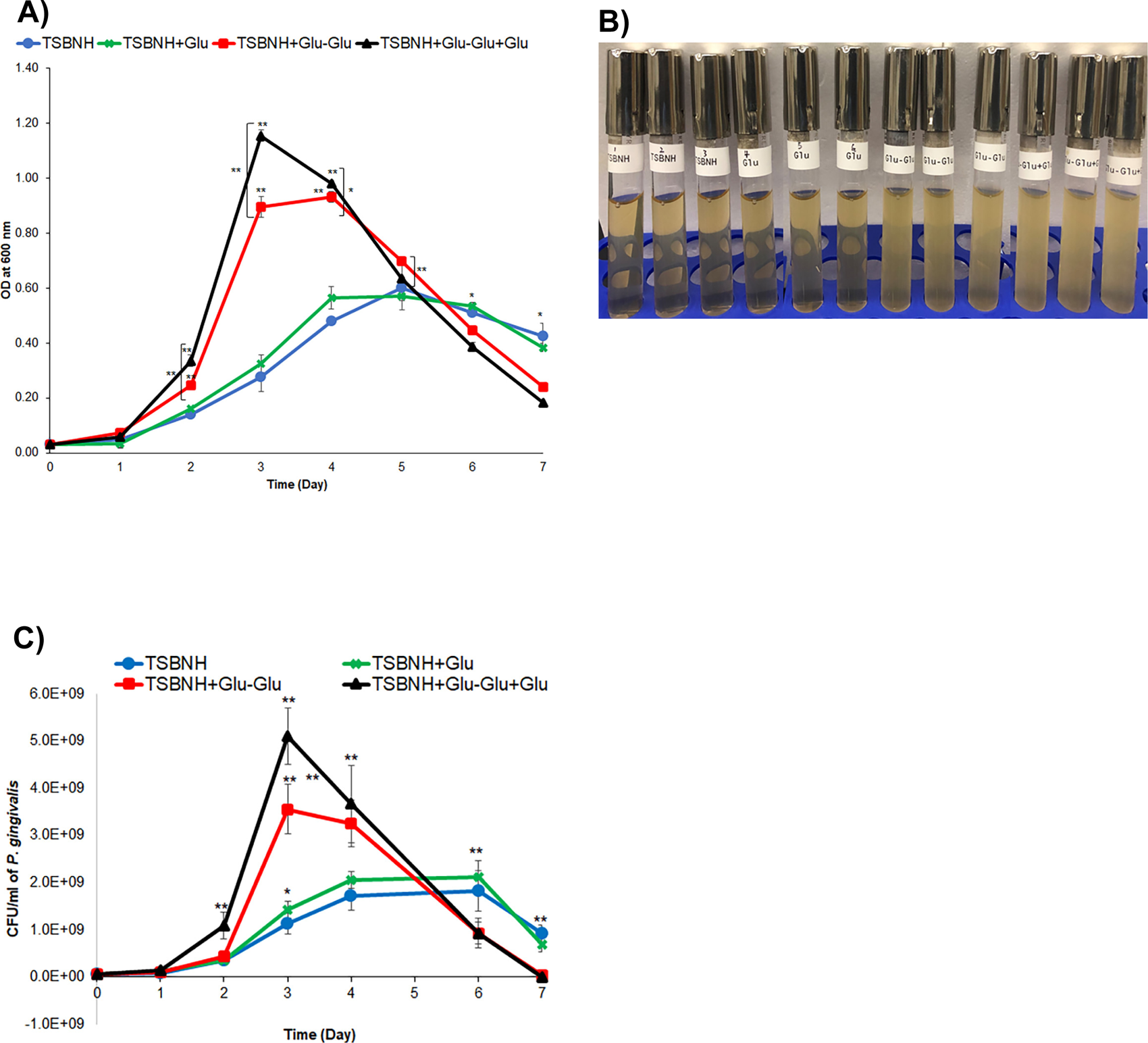
Metabolic activity and growth measurement of wild-type *P. gingivalis* in reduced TSB media (TSBNH) supplemented with Glu-Glu-dipeptide and free glutamate. Wild-type (WT) *P. gingivalis* was grown in TSBNH supplemented with 10 mM Glu-Glu-dipeptide followed by addition of 40 mM free amino acid L-glutamate at Day-1, and the growth was monitored everyday over a 7-day period by measuring optical density (OD) of culture turbidity at 600 (A). Picture of culture tubes were also captured and only the representative picture at Day-3, the time point that showed highest growth (B). The growth of wild-type strain was also monitored by CFU count by drop (10 µl x 3) plating of 10-fold series diluted samples on TSB blood agar (C). The data is represented as Mean ± SD of three replicates. Students’ two-tailed T-TEST was used for statistical analysis and a p-value of <0.05 (*) or <0.01 (**) was considered statistically significant. The assays were repeated at least three times each assayed in triplicates and the results of a representative assay are shown here.

### Atomic Force Microscopy (AFM) imaging showed large cell volume changes of *P. gingivalis* in presence of Glu-Glu dipeptide

Since Glu-Glu dipeptide treatment increased the growth of *P. gingivalis* strain and it is known that bacterial cell volume increases to manage the higher metabolic rate (51), we hypothesized that Glu-Glu-dipeptide treatment will only increase the cell volume of *P. gingivalis* wild-type strain and not the Δ*gdhA* mutant strain. AFM has been successfully used to have 3D ellipsoidal topography of bacterial cells and for the measurement of cell volume from length, width, and height of individual bacterium from taken images (52, 53). Therefore, we performed AFM imaging of *P. gingivalis* wild-type strain grown in either TSB or TSBNH supplemented with Glu-Glu-dipeptide and Δ*gdhA* mutant strain only grown in TSB supplemented with Glu-Glu-dipeptide (as Δ*gdhA* did not grow in TSBNH media) and then estimated the cell volume of *P. gingivalis* strains. The AFM analysis revealed that (**Fig. 10A**) Glu-Glu treatment modulated on the height, length, and width of both *P. gingivalis* wild-type and Δ*gdhA* strains with an inverse relationship. The results showed that Glu-Glu treatment significantly (p<0.01) increased (∼2-fold) the cell volume of *P. gingivalis* wild-type strain in TSB. Similarly, significant (p<0.01) increase in cell volume of wild-type strain was also observed in TSBNH media (**Fig. 10B**). In contrast, the cell volume of *P. gingivalis* Δ*gdhA* mutant strain was significantly less and displayed a lean morphology and the Glu-Glu supplementation further decreased the Δ*gdhA* mutant strain cell volume (become leaner). Although the length of the *P. gingivalis* Δ*gdhA* strain was increased compared to wild-type strain, their width and height were significantly compromised and therefore finally resulted in significantly less cell volume (very lean) compared to the wild-type strain (increased twice in volume) (**Fig. 10B**).

**Fig. 10.**
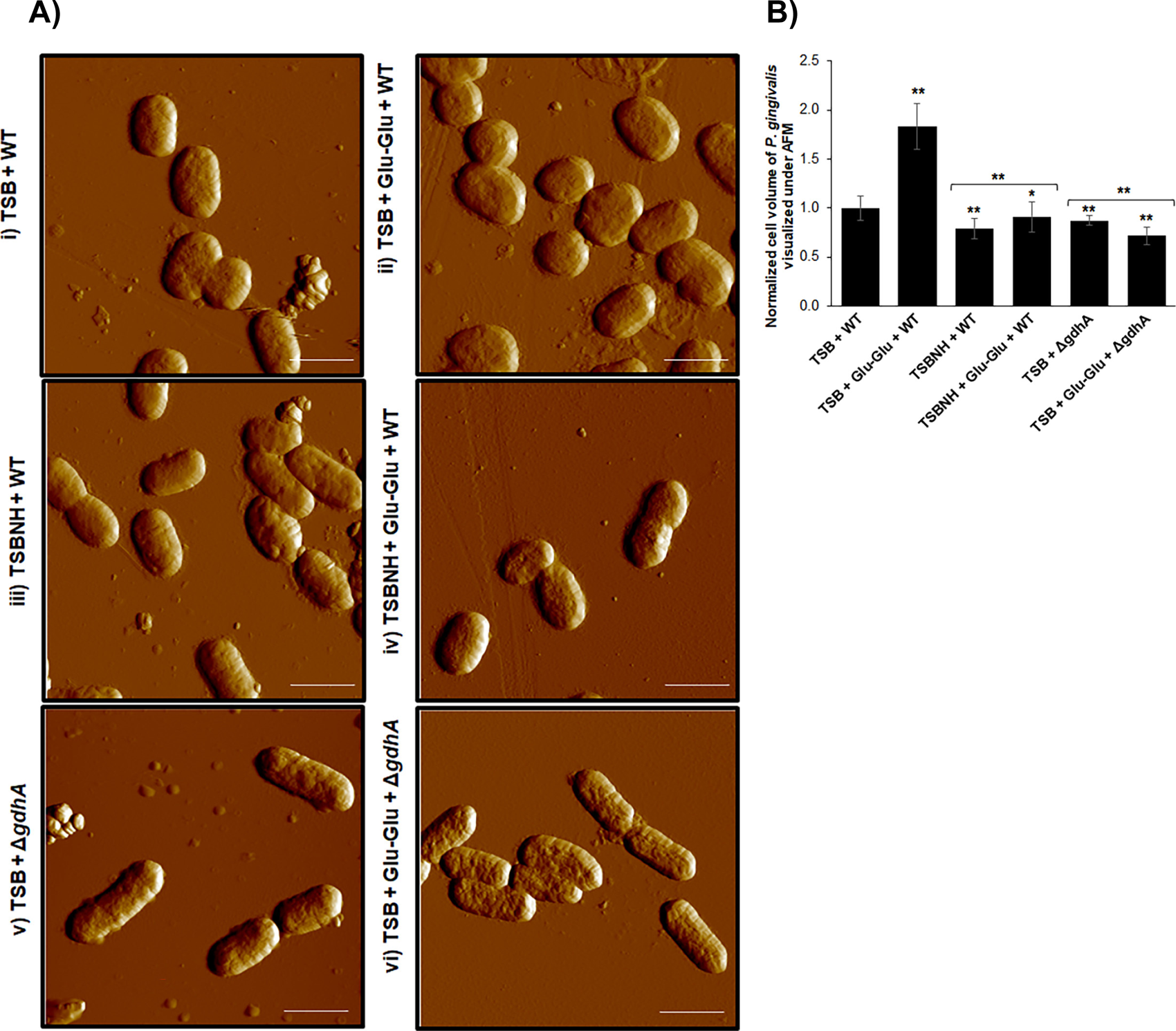
Atomic Force Microcopy (AFM) of *P. gingivalis* strains grown in regular TSB versus reduced TSB (TSBNH) media with/without Glu-Glu dipeptide. Wild-type (WT) and Δ*gdhA* strains of *P. gingivalis* were grown in TSB or TSBNH media supplemented with/without 10 mM Glu-Glu-dipeptideto an early exponential phase (OD_600_ = 0.2-0.3). *P. gingivalis* cells were then harvested by centrifugation and the cells were washed twice with sterile deionized (DI) water and finally resuspended in DI water. A 50 µl bacterial suspension was then dropped on a smooth surface of an AFM-grade mica sheet. Bacterial cells were fixed with methanol and observed under AFM (A). The samples were scanned by contact mode using an AFM probe tip with a stiffness of 0.07 N/m. The assay was repeated at least three times each assayed in duplicates and representative images of each condition are shown here, A (i-vi). The scale bar shown here is 1.0 µM. (B) Cell volume of *P. gingivalis* strains were determined from the measured length, width, and height of at least 25 cells of each condition (see methods and materials for the details). The data is represented as Mean ± SD. Students’ two-tailed T-TEST was used for statistical analysis and a p-value of <0.05 (*) or <0.01 (**) was considered statistically significant.

### Glu metabolism and its key enzyme of *P. gingivalis’* glutamate dehydrogenase (GdhA) are critical for the successful intracellular survival of *P. gingivalis* in human organotypic gingival cultures

Since the organotypic gingival model can mimic the *in vivo* human environment more closely than the human primary GECs for visualization, we have used our novel organotypic gingival culture system (as we recently described in (24) ) to support the importance of Glu metabolism in *P. gingivalis* vacuolar replication and survival for the affluent bacterial colonization in the gingival mucosa. As observed in monolayer primary GECs, the exogenous treatment of organotypic raft with Glu-Glu -dipeptide significantly increased the intracellular level of *P. gingivalis* along with its GdhA levels in the multilayer GECs. In contrast, intracellular *P. gingivalis* level was significantly less in a Δ*gdhA* mutant strain-infected multilayer GECs raft and that was further decreased upon exogenous Glu-Glu treatment (**Fig. 11A, B**). The autophagic marker protein ATG5 which we previously identified to be a central molecule in *P. gingivalis* intracellular survival in GECs (7) displayed both a high degree of induction and colocalization with wild type *P. gingivalis* strain (yellow puncta as a result of colocalization of green fluorescence of *P. gingivalis* and red fluorescence of ATG5) (**Fig. 11C, D**). However, there was a very low level colocalization between the Δ*gdhA* mutant strain and ATG5 (and no increased in level of ATG5 was observed) in the multilayer GEC rafts infected by Δ*gdhA* mutant strain (Data not shown).

**Fig 11.**
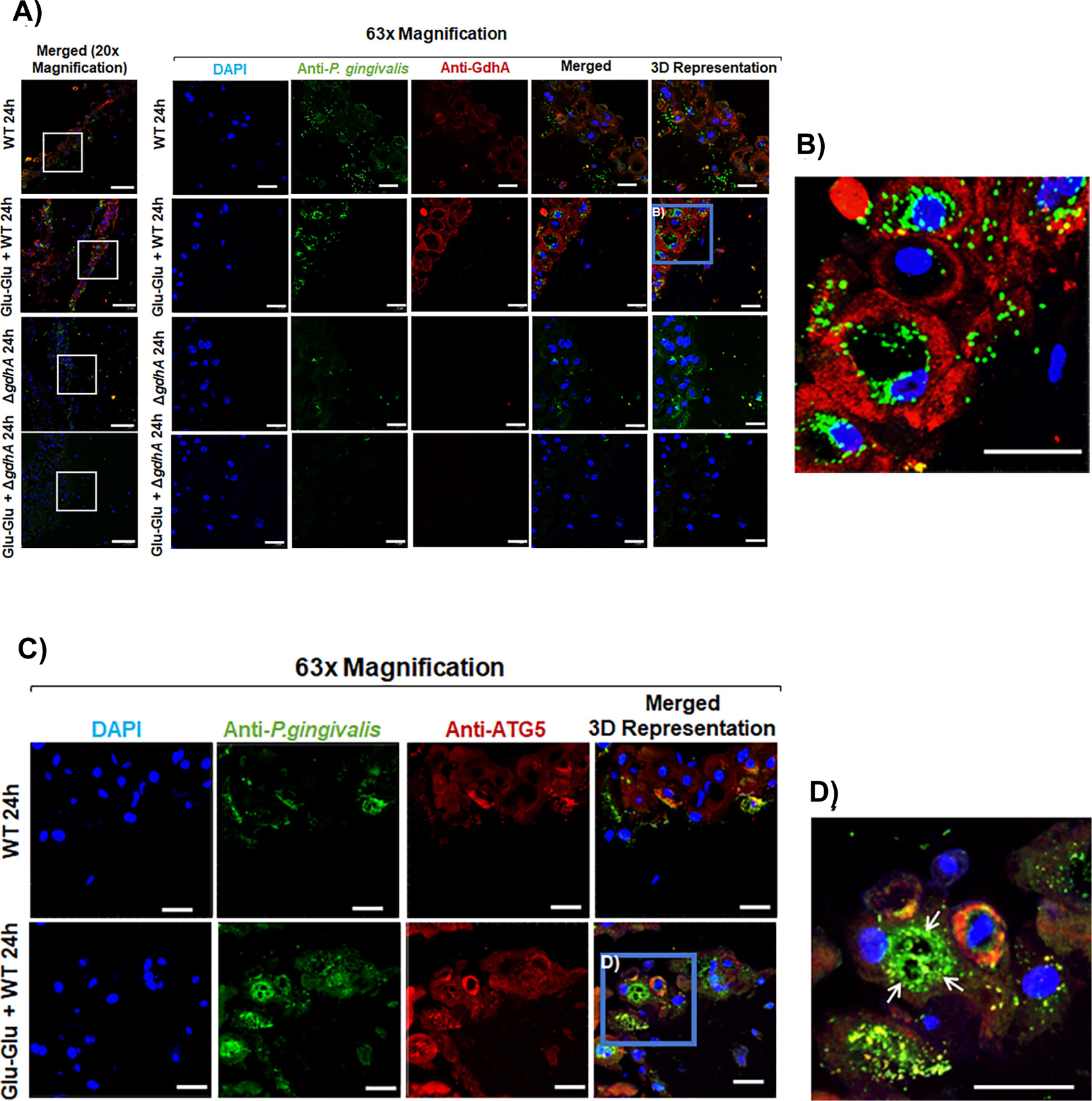
*P. gingivalis’* glutamate dehydrogenase (GdhA) and resulting glutamate metabolism is required for the successful intracellular survival of *P. gingivalis* in Human Organotypic Gingival Cultures. Human primary GECs and Fibroblasts Cells (FBCs) were stepwise co-cultured upon a collagen raft to create an organotypic gingival culture model (as we recently described in (24). Select rafts were then treated with 2 mM Glu-Glu-dipeptide exogenously for 1 h prior to infection Wild-Type (WT) or the isogenic Δ*gdhA* mutant strains (MOI 100) for 24 h. Rafts were then collected and sectioned for immunofluorescence staining. (A) Rafts were stained for *P. gingivalis* (mouse *anti-P. gingivalis*; Alexa 488; green) and GdhA (rabbit anti-rGdhA; Alexa 568; red). (B) A 2x zoomed version of Glu-Glu-dipeptide+ WT. (C) Rafts were stained for *P. gingivalis* (rabbit anti-*P. gingivalis;* Alexa 488; green), ATG5 (mouse anti-ATG5; Alexa 647; purple). (D) A 2x zoomed version of Glu-Glu-dipeptide+ WT indicating enlarged autophagic vacuoles in the cytoplasm of infected GECs (arrows). Samples were imaged using a confocal microscope (Leica DM6 CS Stellaris 5 Confocal/Multiphoton System) at 20x and 63x magnifications. Scale bar is 50 µm for 20x and 20 µm for 63x. Zoomed (2x) and 3D images were obtained via the Imaris software. The range of z-stacks was kept consistent. Representative images of two independent assays are shown here.

Additionally, exogenous treatment of Δ*gdhA* mutant strain-infected raft with Glu-Glu further decreased the levels of Δ*gdhA* mutant strain bacteria. Similarly, there was a low level of colonization by Δ*gdhA* mutant strain to the human organotypic gingiva in general (**Fig. 11A**).

## Discussion

Nutrition uptake and metabolism are critical for all living organisms including bacteria for their survival, reproduction, and persistence. Bacteria has the adaptation capacity to their diverse niches through a variety of molecular mechanisms (46, 54–57) . Additionally, a select number of facultative intracellular bacteria like *P. gingivalis* which have been well adapted to host cellular environments upon environmental perturbations, have evolved the molecular mechanisms to modulate antimicrobial defense systems and exploit homeostatic host metabolic events such as autophagy for survival (4–8, 10). In a nutrition limited environment like autophagic vacuoles, nutrients capturing is a critical challenge for a bacterium such, *P. gingivalis* that is asaccharolytic. We show for the first time here that *P. gingivalis* is highly efficient in uptaking Glu for its metabolism which plays key roles in the vacuolar growth and survival in GECs . The support behind this statement comes from our several lines of strong observations. Our LC-MS/MS analysis of intact *P. gingivalis*-containing vacuoles isolated from *P. gingivalis* infected GECs detects high amount of GdhA of the microorganism and GdhA specific immunostaining results reconfirm the presence of GdhA along with *P. gingivalis* in the vacuoles. The *P. gingivalis* GdhA, like other Gdh homologs, catalyzes the conversion of Glu into α-ketoglutarate, a TCA cycle intermediate. This indicates that *P. gingivalis* might scavenge Glu from the host vacuolar environment for its metabolism. This prediction becomes more prominent when we investigated and found that both whole cell and vacuolar Gln to Glu conversion rate of GECs significantly increases with *P. gingivalis* infection. Intracellular pools of Gln is generally in high levels (36, 58, 59) and Gln can be converted to Glu by glutaminases (38–40).

Our analysis shows that GdhA of *P. gingivalis* does not possess any conventional signal peptide for translocating outside *P. gingivalis* cells while we showed earlier that *P. gingivalis* can secrete an effector without the leaderless motif (4, 60, 61) . Therefore, to explore whether GdhA is secreted from *P. gingivalis,* we superimposed the *P. gingivalis* GdhA structure on GluD, a highly secretory Gdh homolog of *C. difficile,* and that shows significant similarity with their 3D conformations. This indicates that GdhA, like GluD, can be secretory. The biological analysis of bacterial cells free supernatant (CFS) by immunoblot as well as by ELISA confirm that GdhA was highly secreted from *P. gingivalis* cells. Interestingly, both immunoblot, and ELISA results show that supplementation of *P. gingivalis* culture media with Glu-Glu dipeptide, further increases the secretion of GdhA (∼2.5-fold). One of the key indicators of bacterial metabolism is their change in growth/replication. Both our *in situ* antibiotic protection assay and immunofluorescence staining results show significantly higher number of intracellular *P. gingivalis* in GECs with Glu-Glu-dipeptide treatment. In contrast, the GdhA-deficient strain Δ*gdhA* shows significant growth defect and Glu-Glu treatment further impairs the bacterial intracellular proliferation. These findings strongly support the idea that *P. gingivalis* selectively utilizes Glu, and the bacterium specifically induces its GdhA to accomplish this goal. Different intracellular bacteria have different mechanisms to scavenge host nutrients and they have specificity for different nutrients. For example, *Salmonella typhimurium* uptakes intracellular glucose in IL-4-stimulated macrophages for long-term persistence and for that purpose they induce host transcriptional factor PPARδ, peroxisome proliferator-activated receptor, which regulates fatty acid β-oxidation to make more glucose available for the bacterial replication (62). Similarly, chronic infection of *Brucella abortus* and*Mycobacterium tuberculosis* assures glucose availability but by inducing PPARγ, instead of PPARδ (63–66). *Listeria monocytogenes* also induce PPARγ to escape killing by bactericidal factors and inflammatory cytokines ROS, NO, IFNc TNF IL-6 and IL-12 (67). *L. monocytogenes* similarly uptakes glucose from the host and most of the amino acids incorporated in bacterial proteins are derived from intermediates of glucose metabolism (68). *P. gingivalis* cannot metabolize glucose, however, it cannot be ruled out that like *L. monocytogenes, P. gingivalis* may induce host glucose catabolism to increase intracellular amino acids pool especially Glu. Alternatively, *P. gingivalis* may rely on host Gln and convert it to its preferred nutrient Glu by inducing host glutaminases.

Previous studies showed that under planktonic (extracellular) growth condition, *P. gingivalis* can uptake free amino acids like Serine by SstT (PGN_1460) transporter (29, 44). Here we add to the list that *P. gingivalis* can uptake free Glu and this uptake increases over time. But the Glu transporter has yet to be identified. Interestingly, the Glu uptake is significantly higher in a nutritionally reduced TSB media with no hemin (TSBNH) compared to regular TSB media. This finding underlines the preferential use of Glu by *P. gingivalis* during its vacuolar life in GECs, where the nutrient for the microbe is limited. However, uptake of free Glu is not sufficient to support *P. gingivalis* growth unless the growth is first primed with a dipeptide Glu-Glu as we have shown in this study. This is also supported by previous studies where they described that free amino acids alone is not sufficient to augment the growth of *P. gingivalis*, instead it efficiently utilizes dipeptide/oligopeptide (28, 30, 31, 33, 49, 50). Among several dipeptides such as Glu-Glu, Glu-Gln, Gln-Gln, and Arg-Arg tested, Glu-Glu provides the highest growth. A few previous studies also speculated that *P. gingivalis* may consume Glu/Gln containing peptides (30, 31), though none of these reports showed any growth kinetics and/or biological context to directly demonstrate *P. gingivalis* proliferation by Glu metabolism. *P gingivalis* genome analysis shows the presence of multiple dipeptidases (PGN_0250, PGN_0414, PGN_0461, PGN_0788, PGN_1103, PGN_1434, and PGN_1776). Therefore, it is speculated that once dipeptides are transported via proton-dependent oligopeptide transporter, Pot (PGN_0135) (44), RagAB (PGN_0293-0294) (69), and/or oligopeptide transporter Opt (PGN_1518) (44) transporters, dipeptidases work on them to release free amino acids which are then catabolized by specific enzymes like GdhA that specially catabolizes Glu to generate energy and cellular components for *P. gingivalis* growth. To understand the specific role of Glu-Glu in the intracellular growth of *P. gingivalis*, we implemented different *in vitro* approaches to demonstrate that Glu-Glu-dipeptide further increases *P. gingivalis*’ growth in bacterial culture media. Both the culture turbidity in broth and colony forming units (CFUs) on blood agar plates significantly increased over time with Glu-Glu supplementation and it was more prominent when the bacteria was grown in TSBNH media. Although studying growth on agar plate is a widely accepted method to indicate live and culturable bacteria, we reconfirmed the increase of live bacterial number due to Glu-Glu supplementation by an advanced 16S-rRNA-based qPCR method as well (data not shown).

Growth of living organisms depends on their metabolism. As we find significant growth increase in presence of Glu-Glu, we hypothesized that bacterial cells volume should increase for higher metabolism to occur as reported by others previously (52, 53). Interestingly, our AFM analysis of *P. gingivalis* wild-type strain grown in both nutrient rich and reduced media supplemented with Glu-Glu, shows significant increases in cell volume which was opposite in Δ*gdhA* strain. More importantly, Glu-Glu treatment significantly increases the size of *P. gingivalis*-containing autophagic vacuoles from the infected GECs, whereas the Δ*gdhA* mutant strain largely fails to maintain intact autophagic vacuoles with low levels of the Δ*gdhA* mutant strain are detected. Thus, the increase in size of *P. gingivalis*-containing vacuoles may be a vital adaptive strategy by the microorganism to accommodate the space for the new progeny of bacterial cells with potentially enhanced metabolic fitness for protracted survival in GECs (9, 70). These novel findings strengthen our notion that Glu serves as a preferred nutrient for *P. gingivalis* growth especially in an environment where traditional nutrients are limited. Furthermore, it demonstrates for the first time that *P. gingivalis* can control its replication and volume in response to environmental changes, thus assuring adaptation to new niches and conferring increased bacterial metabolic fitness in time under nutritionally fluctuating environments. Intriguingly, a few clinical studies illuminate the clinical significance of Glu in periodontally diseased patient’s oral fluids and serum plasma levels (71, 72). These clinical studies specifically pointed that Glu is highly elevated in saliva of the patients with chronic periodontitis and there are strong positive correlations with the Glu increase and the probing pocket depth, bleeding on probing, clinical attachment level, and plaque index (71, 72). These changes can provide heterogenic superiority to *P. gingivalis* depending on the encountered environments. Moreover, our presented study here which employs sophisticated cellular molecular functional approaches using advanced human primary GEC-based systems combined with the rigorous bacterial genetics presents a pivotally novel understanding on *P. gingivalis*’ growth, survival requirements and behavior in the autophagic vacuoles (**Figs. 11 and 12**). Since the therapeutic targeting of intracellular bacteria stands as a formidable challenge in the periodontal disease treatment where antimicrobials first need to effectively pass-through host cells phospholipid bilayer before reaching to the bacteria and it becomes even more difficult to reach bacteria like *P. gingivalis* that mainly shelters inside the highly complex double membrane vacuoles with a variety of autophagy proteins tightly integrated (7, 24). This belief further corroborates to the clinically recalcitrant nature of *P. gingivalis* to conventional periodontal therapies such as scaling and root planing combined with various antimicrobial adjuvants (20, 22) . Therefore, our results underscore that Glu metabolic pathways could be novel targets for therapeutic interventions to control *P. gingivalis* chronic presence in the oral mucosa which act as crucial reservoir for *P. gingivalis* systemic dissemination and dysbiosis.

**Fig. 12.**
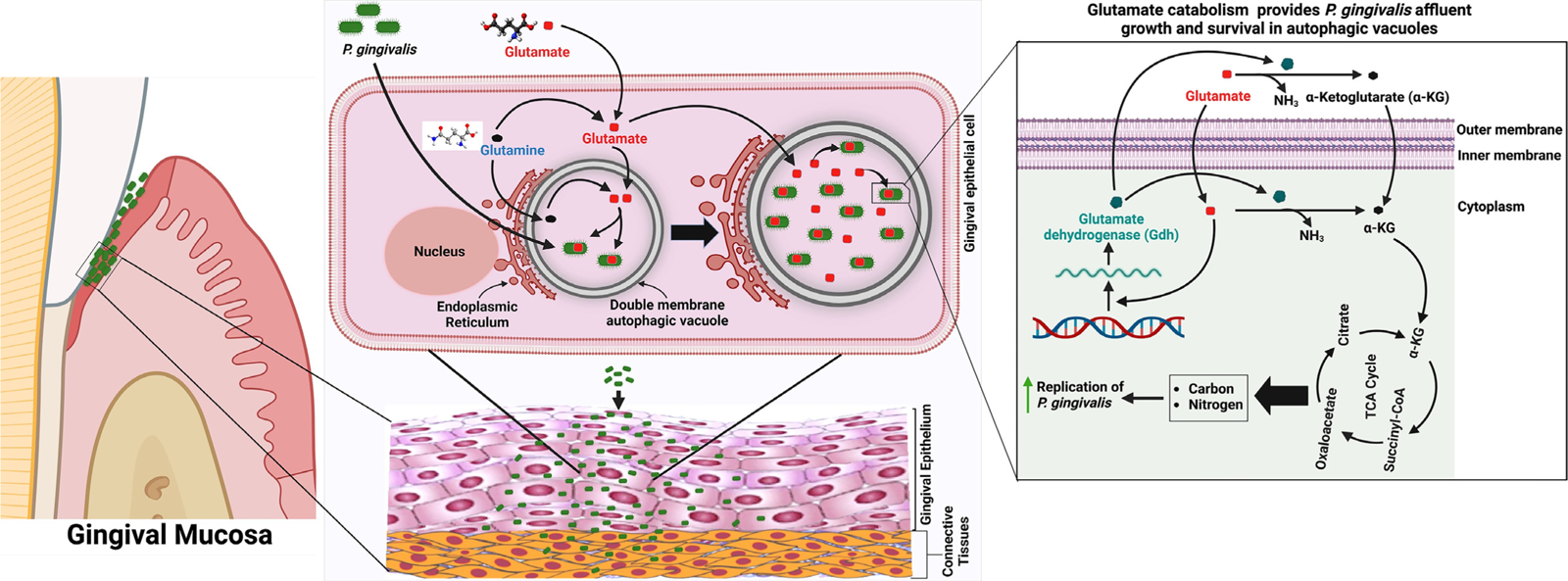
Schematic representation on the proposed impact of glutamate catabolism on *P. gingivalis* growth and survival in the GEC intracellular vacuolar environment and successful colonization in the oral mucosa. *P. gingivalis* invades human GECs of gingival mucosa and traffics into ER-rich double membrane autophagic vacuoles (7, 24). *P. gingivalis* infection favors **i**ntracellular conversion of Gln to Glu (endogenous Glu). Exogenous Glu supplementation along with endogenous Glu serve as preferred nutrient for *P. gingivalis* for their proliferation and survival in the intracellular vacuoles in GECs. Glu catabolism also significantly increased the size of the vacuoles to facilitate more space for new progeny of bacteria. Glutamate dehydrogenase (GdhA), which is secreted in vacuoles from *P. gingivalis* plays a key role in Glu catabolism. GdhA converts Glu to α-ketoglutarate which is then shuttled into TCA cycle. Glu metabolism provides carbons and nitrogen for anabolism of macromolecules necessary for organismal growth. The chronic presence of *P. gingivalis* as has been found in severe periodontitis gingival tissues may spread the infection to the deeper connective tissues and lead to the systemic infections (20, 24) .

## Materials and Methods

### Bacterial and human primary gingival epithelial cells culture

*P. gingivalis* ATCC 33277 and its isogenic glutamate dehydrogenase gene *(gdhA,* encoded by *PGN_1367*)[deletion mutant (Δ*gdhA*) strains were cultured to early exponential phase (OD_600_ ∼ 02.-0.4) in trypticase soy broth or TSB (BD, Sparks, MD) supplemented with yeast extract (BD) (1 mg/mL), menadione (Chem-Implex Int’l Inc., Wood Dale, IL) (1 μg/mL), and hemin (Millipore-Sigma, St. Louis, MO) (5 μg/mL) at 37 °C under anaerobic conditions and harvested as previously described (5, 10, 73, 74). Here after this media composition was termed as regular TSB media. A nutritionally limited TSB media was prepared with same composition as TSB but lacks hemin (no added hemin) and named as TSBNH (TSB with <u>N</u>o <u>H</u>emin). Erythromycin (10 µg/mL, Fisher Scientific, Pittsburgh, PA) was added to the TSB for selective growth of the Δ*gdhA* strain. The cells pellet of *P. gingivalis* were resuspended in Hyclone Dulbecco’s phosphate[buffered saline (DPBS) containing calcium and magnesium (GE Healthcare, Pittsburgh, PA), and were quantified using a Klett[Summerson photometer (Bel-Art, Wayne, NJ). Human primary gingival epithelial cells (GECs) were obtained and cultured as described previously (10, 74, 75). Human primary gingival epithelial cells (GECs) were obtained following the oral surgery of adult patients who were accepted for either tooth crown lengthening procedures or impacted third molar extraction. Following the randomization and anonymization of the patients, under the approved guidance of the University of Florida Health Science Center Institutional Review Board (IRB, human subjects assurance number FWA 00005790), healthy gingival tissue samples were collected. Cells were cultured in a serum[free keratinocyte basal medium or KBM (Lonza Inc., Walkersville, MD) without any antibiotics at 37 °C in 5% CO2. GECs were cultured at 70%-80% confluence and infected with *P. gingivalis* strains at a multiplicity of infection (MOI) of 100 whenever appropriate.

### Proteomic analysis of autophagic vacuolar samples from human GECs infected with *P. gingivalis*

*P. gingivalis* wild-type strain ATCC 33277 was grown to an early exponential phase (OD_600_ ≈ 0.2-0.4) and following centrifugation at 10,000 x *g* for 5 min, cells pellet was resuspended in sterile DPBS. It was then incubated in 5 μg/mL Lipobiotin in PBS overnight at 4°C. Labeled *P. gingivalis* was then incubated with MagCellect Streptavidin Ferrofluid (R&D Systems, Inc., Minneapolis, MN) at a concentration of 7– 14[×[10^8^ particles per 2[×[10^8^ bacteria for 20 min at 4°C. To isolate *P. gingivalis*-containing intact autophagic vacuoles, GECs were plated at a density of 1.5 × 10^6^ cells per 100 mM dishes and infected with wild-type *P. gingivalis* at MOI 100 for 3 h, 6 h, and 24 h, and *P. gingivalis*-containing vacuoles were then isolated following a special protocol of Steinhauser *et al* (76) adapted for *P. gingivalis*. In brief, GECs were washed with buffer A (15[mm HEPES buffer with 20[mm sucrose, 50[mm MgCl_2_, and protease inhibitors containing 0.02% ethylenediaminetetraacetic acid) to remove non-ingested bacteria, were trypisinized, and resuspended in buffer A combined with benzonase nuclease (25 U/mL; Millipore-Sigma) and Cytochalasin D (5 µg/mL; Millipore-Sigma). Following resuspension, cell membranes were disrupted, and *P. gingivalis*-containing vacuoles were released via sonicating the samples on ice via a ThermoFisher Qsonica Sonicator Q500 (FisherSci) at an amplitude of 75% for 10 s. Samples were then centrifuged at 300 x *g* for 2 min between sonication steps, after which the supernatant was removed and replaced with 500 μL of buffer A. After the third sonication, the samples were left on ice for 10 min. The supernatants were then removed and incubated at 37°C for 5[min. Following incubation, the samples were placed in a Magnetic Rack (Bio-Rad) for 30 min prior to repeated manual rinsing of the magnetic fraction with buffer A at a flow rate of 250 μL/min for 15[min to ensure purity. Finally, the vacuoles-containing fractions were collected and centrifuged at 15,000 x *g* for 12 min, before the supernatant was decanted and the vacuoles were resuspended in 100 μL of HEPES buffer combined with protease inhibitors. The autophagic vacuoles were used for label-free quantification (LFQ) of *P. gingivalis*’ proteins enriched in it by LC-MS/MS using Orbitrap Fusion Lumos with ETD/UVPD, a high resolution and high mass accuracy instrument (Thermo Fisher Scientific) available at Proteomics core, Medical University of South Carolina. Level of GdhA (PGN_1367, Protein ID # B2RKJ1) was quantified based on the peak intensity of the corresponding MS spectrum of GdhA with 95% confidence. GdhA was confirmed by BlastP searches using individual peptide fragments as well as all fragments together. The assay was repeated two times and at least six replicates were pooled together in each assay.

### Immunofluorescence staining of autophagic vacuoles for GdhA and *P. gingivalis*

As mentioned above, GECs seeded on 100 mm dishes and pre-treated with 2 mM Glu-Glu (EE) for 1 h prior to infection with wild-type or Δ*gdhA* strains of *P. gingivalis* at MOI 100 for 6 h. Vacuoles were then isolated following the same protocol mentioned above and 10-20 µL of isolated vacuoles were placed on a microscope slide and fixed with 10% neutral buffer formalin for 30 min at room temperature. The samples were then blocked with 3% BSA for 30 min and stained for GdhA and *P. gingivalis* following standard protocol (5–9). In brief, fixed samples were incubated with primary antibodies of rabbit anti-rGdhA (rPGN_1367) or anti-mouse *P. gingivalis* 33277 (1∶1000) for 1 h at room temperature. Samples were washed three times and incubated with donkey anti-rabbit Alexa fluor 594-conjugated secondary antibody (1∶2000, Invitrogen, Carlsbad, CA; Cat. # R37119) or goat anti-mouse Alexa fluor 488-conjugated secondary antibody (1∶2000, Invitrogen, Carlsbad, CA, Cat. # A-10680) for 1 h to stain GdhA as red or *P. gingivalis* as green. To check the integrity of vacuoles, samples were stained with thiol-tracker violet (Invitrogen # T10095) as it stains the thiol (sulfhydryl) groups present on vacuoles (77) before mounting with anti-fade mounting media. The vacuoles were then imaged by a confocal microscope (Leica DM6 CS Stellaris 5 Confocal/Multiphoton System) at 63x magnification where the scale bar is 5 µm. Images were analyzed by Imaris Microscopy imaging software (Oxford Instrument). The diameter (µm) of representative vacuoles were measured. Additionally, the surface area of at least 25 vacuoles from three independent assays was also quantified and normalized to *P. gingivalis* wild-type (WT) 6 h infection. The data is represented as Mean ± SD and Students’ two-tailed T-TEST was used for statistical analysis where a p-value of <0.05 (*) or <0.01 (**) was considered as statistically significant.

### Intracellular whole cell and vacuolar conversion of glutamine to glutamate by *P. gingivalis*

GECs were seeded in either 96-well plates (1.0 x 10^4^ cells/well) for intracellular or 100 mm dishes (1.5 x 10^6^ cells/plate) for intra-subcellular (i.e., vacuolar) conversion of glutamine (Gln) to glutamate (Glu) assay using Glutamine/Glutamate-Glo Assay kit (Promega, Madison, WI; Cat. # J8021) following kit’s protocol. In brief, GECs were pretreated for 1 h with 2 µM exogenous Gln and infected with *P. gingivalis* wild-type strain for 6 h or 24h. Cells were washed with PBS after removing the media and stopped the endogenous reaction and lysed the cells with inactivation solution I (0.3 N HCl in 450 mM Tris, pH8.0). For intracellular in GECs, cell lysates were used for the assay. For intra-subcellular, intact *P. gingivalis* containing autophagic vacuoles were selectively isolated using the protocol (76) mentioned above and the isolated vacuoles were used for the assay. The luminescent signal proportional to the amount of glutamate was recorded by a Biotek H1M monochromatic plate reader. Glu concentrations with and without exposure to kit-provided glutaminase treatment were obtained and the relative amounts of Glu were calculated by comparison with the standard curve prepared with known concentrations of glutamate. The assay was repeated two times with each assayed in six replicates.

### Glutamate binding prediction with GdhA of *P. gingivalis* by molecular docking

For molecular simulation of potential Glu binding to GdhA of *P. gingivalis*, 3D structure of Glu molecule was created by Avogadro software and saved as PDB file and the 3D structure of GdhA (Protein ID# B2RKJ1) was extracted from UniProt website. The PDB files of GdhA and Glu were submitted for protein and ligand molecular docking model generation by supercomputer at ClusPro (78–81). The balanced model having the best cluster size was chosen for analysis by PyMOL. The Glu binding sites in GdhA are shown in purple color, active sites in Red color, and its cofactor NAD+ binding sites in Blue color. Since glutamate dehydrogenase (GluD) of *Clostridium difficile* is highly secretory (47) and the *GdhA* of *P. gingivalis* was secreted at high intensities inside the vacuoles, we superimposed their 3D structures using PyMOL (SchrÖdinger, LLC; New York) molecular docking visualization system. PDB files of 3D structure of GluD (Protein ID# P27346) were also obtained from <u>UniProt</u> website. GdhA and GluD were superimposed by PyMOL tool. The cartoons of GluD and GdhA were colored red and green, respectively. FASTA sequences of Glud and GdhA were also aligned using Constraint-based Multiple Alignment Tool or COBALT (82) available via NCBI BlastP program. The alignment was set as compact view to identity conservation parameter where red color indicates highly conserved residues and blue indicates less conserved ones.

### Confirmation of GdhA secretion from *P. gingivalis* in bacterial cell free supernatant (CFS) by Immunoblot

To validate the *in silico* prediction of GdhA secretion form *P. gingivalis*, CFS prepared from *P. gingivalis* wild-type (WT) or Δ*gdhA* strains were immunoblotted with rGdhA-specific antibody prepared in this study (see below). In brief, a 200 mL of *P. gingivalis* wild-type (WT) strain with/without 2 mM Glu-Glu (EE) dipeptide and a 400 mL of isogenic *gdhA* deletion mutant (Δ*gdhA)* strain was grown to mid exponential phase (OD at 600 nm were about 0.9 for WT strain and 0.45 for the Δ*gdhA* strain). Bacterial cells were removed by centrifugation at 10,000 x g for 10 min at 4 °C. The supernatant was filtered (0.1 µm pore size) to make it cell free supernatant (CFS) and proteins were precipitated by salting-out (83) (Duong-Ly and Gabelli SB, 2014) with (NH_4_)_2_SO_4_ (about 80% saturation) followed by centrifugation at 20,000 x g for 1 h at 4 °C. The precipitate was resuspended in 5 mL of cell lysis buffer (20 mM Tris-Cl, pH 8.0; 20% glucose, 1 mM EDTA, 0.1 mM DTT, and 1 mM PMSF) little modified from (84). The samples were dialyzed (Fisher Scientific # 21-152-10) against 400x excess PBS buffer at 4 °C overnight with two changes of the PBS buffer. The samples were then concentrated using Amicon ultra 10-kDa cutoff filter unit (Millipore-Sigma # UFC9010) to a 1 mL. The concentrated CFS were collected in a 1.5 mL Eppendorf tube and briefly centrifuged at 10,000 x g for 3 min to remove any residual slats and debris. The collected supernatants were used to estimate protein concentration by Bradford assay. A 40 µg of protein of CFS samples and 10 ng of purified rGdhA were resolved in a 4-20% SDS-PAGE (Bio-Rad, Hercules, CA; # 4561094) and immunoblotted with affinity purified anti-rabbit rGdhA antibody (0.3 µg/mL, GenScript, Piscataway, NJ). Image J software was used for densitometric analysis of the blots for GdhA, where data is represented as Mean with standard deviation of two replicates and students’ two-tailed T-TEST was used for statistical analysis. A p-value of <0.01 (**) was considered Statistically significant. Note that a 10 µL of this CFS was drop plated on TSB blood agar or in TSB to confirm if there was presence of any live *P. gingivalis* cells. Additionally, 2 µL CFS was used in 25 µL reaction volume for *P. gingivalis* 16S rRNA-specific PCR to see if there was any DNA of *P. gingivalis* that might come from lysed *P. gingivalis* cells. The presence of DNA could void the claim that GdhA was secreted in the CFS, instead it would indicate that GdhA was leaked into the CFS from lysed bacterial cells.

### Reconfirmation and quantification of GdhA in CFS by ELISA

To further confirm the presence and better quantify the level of GdhA in CFS, the CFS samples were analyzed by indirect ELISA. A 100 µL of 10-fold series dilutions of CFS samples of WT strain (with/without 2 mM Glu-Glu) and Δ*gdhA* strains with plate coating buffer (R & D Systems, Minneapolis, MN; # DY006) were used to coat a 96-well plate (R & D Systems, # DY990) overnight at 4 °C. Similarly, 100 µL of different concentrations (0 to 20 ng/mL) of purified rGdhA were used to coat the plate to prepare standard curve. The coated plate was washed once with washing buffer (0.05% tween-20 in PBS buffer, i.e., PBST) before blocking with blocking buffer (1%BSA in washing buffer) for 1 h at 37 °C. The plate was washed once with washing buffer before incubation with 100 µL of Rabbit anti-rGdhA antibody (0.05 µg/mL in blocking buffer) overnight at 4 °C. The plate was washed 4-times with washing buffer and remaining drops were removed by patting the plate on paper towel. A 100 µL (1:20,000 dilution) of Mouse Anti-Rabbit IgG Fc monoclonal antibody (mAb, 14H9H10) conjugated with HRP (GenScript # A01856) was added to each well and incubated at 37 °C for 30 min. This mAb reacts only with the Fc portion of rabbit IgG but not with the Fab portion of rabbit IgG and has no cross-reactivity with immunoglobulins of other species tested (GenScript). Since polyclonal antibodies can recognize many epitopes, the background and the cross reactivity in assays is usually high, use of this mAb minimizes the background and cross reactivity (GenScript). The plate was then washed with washing buffer 4-times and 100 µL of TMB substrate reagent (Thermo Fisher Scientific # N301) was added into each well and incubated at room temperature. After sufficient color development (20 min), a 100 µL of stop solution (0.16 M H_2_SO_4_, Thermo Fisher Scientific # N600) was added and the absorbance was recorded at 450 nm using a Biotek H1M monochromatic plate reader. The amount of GdhA in the samples (accepted absorbance values were the linear range of the standard curve) were calculated from the formula obtained from the standard curve. The values with no rGdhA were used as Blank that Blank values were subtracted from each sample and standards. To reconfirm the specificity of the primary rGdhA antibody, a high concentration (5 µg/mL) of BSA was used. The assay was repeated twice each assayed in triplicates. The data is represented as mean with standard deviation and a P-value of <0.01 (**) was considered as Statistically significant which was determined by Students’ two-tailed T-TEST.

### Glutamate dehydrogenase (Gdh) activity assay

The Glutamine/Glutamate-Glo assay kit as mentioned above was modified to measure the activity of Gdh. The principle of the kit is that Gdh utilizes glutamate and NAD^+^ to generate α-ketoglutarate and NADH. A reductase enzyme with the help of NADH then converts a pro-luciferin reductase substrate to luciferin which is then utilized by the Ultra-Glo recombinant luciferase to produce light (luminescence). The luminescent signal is proportional to the amount of glutamate consumed. Based on this principle, the Gdh activity in the CFS of *P. gingivalis* WT (with/without 2 mM EE) and Δ*gdhA* strains and 1.0 μg of purified rGdh were used as the Gdh source in the reactions for the unknown samples. The kit provided Gdh was used as positive control and reaction mix without any Gdh was used as Blank. For the standard curve preparation, different concentration (kit recommended) of glutamate was used. The assay was modified based on empirical analysis to increase the glutamate (substrate) concentration to 20 mM and NAD^+^ concentration to 3 mM for Gdh activity assay in CFS samples. The luminescence was recorded every 5 min interval over 12 h by a Biotek H1M plate reader. The relative light unit or RLU (luminescent) was plotted against different standard concentrations of glutamate to prepare the standard curve. For the samples, the whole RLU data was first plotted against all the time (min) points and then RLU values in the linear range of the standard curve were used to determine the Gdh activity (nmol/min/μg protein) using the following formula: ([Glu_final_-Glu_initial_] x 1000)/([T_final_-T_initial_] x μg protein), where Glu_final_ and Glu_intial_ are amount of glutamate catabolized at time T_final_ and T_intial,_ respectively, calculated from the standard curve using the respective RLU values of the samples. Here, 1000 was the multiplication factor to convert μmole to nmole. The amount of total protein present in the CFS was determined by Bradford assay. The assay was repeated twice each assay in duplicates. The data is represented as mean with standard deviation and a P-value of <0.01 (**) was considered as Statistically significant which was determined by Students’ two-tailed T-TEST.

### Construction of Δ*gdhA* (PGN_1367) deletion mutant strain in *P. gingivalis* ATCC 33277 strain

In-frame deletion of *gdhA* gene (1338 bp, 445 aa) was performed by replacing entire *gdhA* gene except start codon ATG and stop codon TAA (i.e., 1332 bp was deleted) with the *ermF* gene (801 bp, 266 aa) that confers resistance to erythromycin. The *ermF* gene with its own start and stop codons was inserted in between the start and stop codons of *gdhA* gene. Therefore, the *gdhA* locus in Δ*gdhA* strain was (1332-801=531) 531 bp shorter than the wild-type *gdhA* locus. A standard overlapping PCR technique (10, 54, 85) was used with little modification. Briefly, about 1000 bp upstream and downstream flanking regions of *gdhA* gene from *P. gingivalis* wild-type strain ATCC 33277 genomic DNA and *ermF* gene from pPR-UF1(86) were PCR amplified using the primers listed in Table 1. Q5 high-fidelity Taq DNA polymerase (New England Biolabs Inc., Ipswich, MA) which has a 3’→ 5’ exonuclease activity and hence results in ultra-low error rates was used for PCR. Primers were designed to have a 40 bp overlapping region along with the target sequence. A one-step cloning of three PCR fragments putting *ermF* gene in the middle of upstream and downstream flanking fragments of *gdhA* gene was used to assemble them into pJET1.2/blunt vector (Thermo Fisher Scientific, Grand Island, NY) using RepliQa HiFi assembly kit (Quantabio, Beverly, MA) and transformed into *Escherichia coli* DH5α. The recombinant plasmid DNA was used to PCR amplify the insert using pJET1.2-F and pJET1.2-R primers (Table 1), and 1.0 μg of purified DNA was used to electroporate freshly prepared competent cells of wild-type *P. gingivalis* following standard protocol(86). The mutant strain was selected on TSB medium supplemented 1.5% Bactro agar (BD), 5% sheep blood (Hemostat Laboratories, Dixon, CA), and 10 µg/mL of erythromycin(10, 86). The mutant was confirmed by PCR using flanking primers (PGN_1367F and PGN_1367R, Table 1) designed from 50-100 bp away from the upstream and downstream fragments, as well as by sequencing. All primers were purchased from Eurofins Genomics LLC (Louisville, KY) and PCR product sequencing also performed by Eurofins.

**Table 1.**
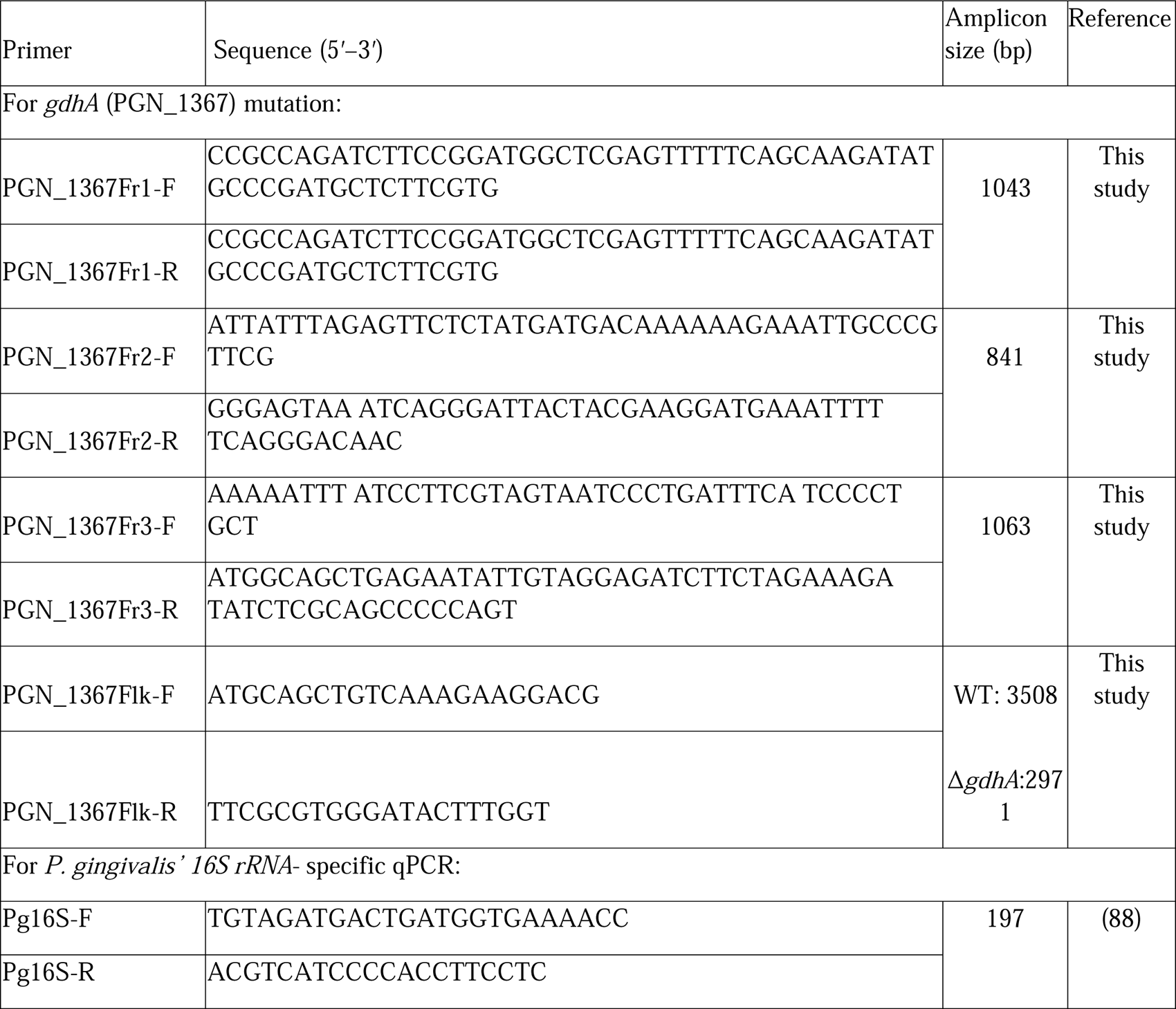
Primers used in this study.

### Antibiotic protection assay and Immunofluorescence (IF) staining for comparative intracellular level quantification of *P. gingivalis* wild-type and Δ*gdhA* strains in human GECs

To investigate effect of Glu on comparative intracellular survival and replication of *P. gingivalis* wild-type and its isogenic Δ*gdhA* strains in human GECs, standard antibiotic protection assay and IF staining were performed (4–10, 35, 73, 87). For antibiotics protection assay, GECs were seeded at a density of 2 x 10^5^ cells/well of six-well plates (Fisher Scientific# 07-200-83). Cells were pre-treated with 2 mM Glu-Glu for 1 h prior to infection with wild-type or Δ*gdhA* strains of *P. gingivalis* at MOI 100 for 3 h, 6 h, and 24 h. Cells were washed 3 times with PBS and treated with gentamicin (300 µg/mL) and metronidazole (200 µg/mL) for 1 h. Media was aspirated and cells were washed with PBS to remove any residual antibiotics and debris before collection in 1 mL of Qiazol reagent (Fisher Scientific# NC9055817). RNA was isolated following the Qiazol reagent procedural guidelines. DNA was removed from the RNA with the Ambion Turbo DNA-free kit (Invitrogen). cDNA was synthesized from 1 µg of total RNA using high-capacity cDNA reverse transcription kit (Applied Biosystems, Foster City, CA). A 1:10 diluted cDNA was used to detect *P. gingivalis* by qRT-PCR in CFX96 real-time system (Bio-Rad, Hercules, CA) using *P. gingivalis* 16S rRNA-specific primers(88) and conditions used before(6, 73). The CFUs was calculated from the Ct (Threshold cycle) value of each sample with the help of a standard curve which was prepared with serial dilutions of genomic DNA of *P. gingivalis* of known CFU and their corresponding Ct values(73, 89).

For IF staining, GECs (8 x 10^4^ cells) were seeded in 4-well chamber slides (Fisher Scientific# 12-565-2) and pre-treated with 2 mM Glu-Glu for 1 h prior to infection with wild-type or Δ*gdhA* strains of *P. gingivalis* at MOI 100 for 3 h, 6 h, and 24 h. The cells were then washed several times with PBS before fixation with 10% normal buffered formalin. The cells were then permeabilized with 0.1% Triton X-100, blocked with 3% BSA for 30 min. Anti-rabbit *P. gingivalis-*specific antibody (1:1000) and a goat anti-rabbit Alexa fluor 488-conjugated secondary antibody (1∶2000, Invitrogen, Carlsbad, CA; Cat. # A27034) was used to stain *P. gingivalis* as green. Rhodamine Phalloidin was used to stain the cells cytoskeleton as red. The samples were washed and mounted with coverslips using VECTASHIELD vibrance antifade mounting medium with DAPI (Vector Laboratories #H-1800) to stain cells nuclei as blue and visualized under a confocal microscope (Leica DM6 CS Stellaris 5 Confocal/Multiphoton System) at 20x/63x magnification). At least two separate experiments each assayed in duplicates were performed. The scale bar is 100 µm for 20x Magnification and 20 µm for 63x Magnification. The mean intensity of green fluorescence of at least 25 cells of each indicated condition derived from intracellular *P. gingivalis* was quantified by ImageJ software and the data is presented as Mean ± SD. Students’ two-tailed T-TEST was used for statistical analysis and a p-value of <0.01 was considered statistically significant.

### Glutamate uptake by *P. gingivalis* and preference to Glu-Glu dipeptide

*P. gingivalis* wild-type strain was incubated in 5 mL of TSB or TSBNH (a nutritionally limited TSB media that lacks hemin) media with or without radiolabeled L-[^14^C(U)] Glutamate (1.0 mCi/mmol, Perkin Elmer # NEC290E050UC) at 37°C under anaerobic condition. Bacterial growth was monitored by measuring OD at 600 nm and 1 mL of the culture was collected at 1 h, 18 h, and 48 h time points to harvest bacterial cells by centrifugation at 10,000 x *g* for 5 min. Bacterial cells pellet was washed three times with PBS to remove any glutamate adhered outside of the cells. Cells were then lysed with 20 mg/ mL of lysozyme (Millipore-Sigma, # 4403) and mixed with 5 mL of liquid scintillation cocktail (Beckman Coulter # 141349, Brea, CA) before measuring the radioactivity (Counts per minute [CPM]) using a liquid scintillation counter LS 6500 (Beckman Coulter). Bacterial culture without any radiolabeled glutamate was used as blank and the value was subtracted from the test samples. Glutamate uptake by *P. gingivalis* was normalized to OD_600_ of *P. gingivalis* growth and, hence, expressed as CPM/OD_600_. The assay was repeated twice with each assay in triplicates. The data is represented as Mean ± SD, where n = 3; and a p-value of <0.01 was considered statistically significant (Students’ two-tailed T-Test). Since *P. gingivalis* showed significant more uptake of ^14^CU-Glutamate in TSBNH (a nutritionally limited) media, TSBNH media was used to test growth of *P. gingivalis* in different dipeptides Glu-Glu (Millipore-Sigma), Glu-Gln, Gln-Gln, or Arg-Arg (later three were synthesized from RS Synthesis LLC, Louisville, KY). *P. gingivalis* wild-type strain was grown in a 10 mL TSB for 48 h. Culture was then centrifuged at 10,000 x *g* for 5 min and cells pellet was resuspended in TSBNH to adjust OD_600_ of about 4. The culture was then diluted to OD_600_ of 0.03 in 7 mL of TSBNH supplemented with 10 mM dipeptide. The growth was monitored everyday over a 5-day period by measuring optical density (OD) of culture turbidity at 600 nm. The data is represented as Mean ± SD of three replicates. Students’ two-tailed T-TEST was used for statistical analysis and a p-value of <0.05 (*) or <0.01 (**) was considered statistically significant. The assay was repeated at least three times.

### Growth assay of *P. gingivalis* wild-type strain in TSB media or in TSBNH media with Glutamate precursors

Since *P. gingivalis* depend on hemin from the environment for their growth and iron (present in hemin) metabolism provides this microorganism their characteristic black pigment(90–93), TSB media was modified to make a nutritionally reduced TSB media, TSBNH to see if complete removal of hemin from TSB media would be effective to capture better growth difference with/without supplementation of Glu-Glu dipeptide as glutamate precursor. *P. gingivalis* wild-type strain was grown in a 10 mL TSB for 48 h. Culture was then centrifuged at 10,000 x *g* for 5 min and cells pellet was resuspended in TSBNH to adjust OD_600_ of about 4. The culture was then diluted to OD_600_ of 0.03 in 7 mL TSB, or TSBNH supplemented with 10 mM Glu-Glu dipeptide (Millipore-Sigma). The growth was monitored everyday (unless specified) over a 7-day period by measuring optical density (OD) of culture turbidity at 600 nm. The data is represented as Mean ± SD of three replicates. Students’ two-tailed T-TEST was used for statistical analysis and a p-value of <0.05 (*) or <0.01 (**) was considered statistically significant. The assay was repeated at least three times.

### Growth assay of *P. gingivalis* wild-type or Δ*gdhA* strains in TSB or TSBNH media supplemented with Glutamate precursors

*P. gingivalis* wild-type strain was grown in a 10 mL TSB media for 48 h and processed similarly as mentioned above. But for Δ*gdhA* strain, it was grown in a 100 mL TSB with 10 µg/mL erythromycin for 72 h. Culture was centrifuged at 10,000 x *g* for 5 min and cells pellet was resuspended in TSBNH to adjust OD_600_ of about 4. The culture was then diluted to OD_600_ of 0.03 in either 7 mL TSB or TSBNH supplemented with 10 mM Glu-Glu dipeptide (Millipore-Sigma). A 40 mM free amino acid L-glutamate (E) (TCI America # G0188, Portland, OR) was added after 6 h (wild-type) or 24 h (Δ*gdhA*) of growth whenever appropriate. The growth was monitored everyday over a 7-day period by measuring optical density (OD) of culture turbidity at 600. Simultaneously, 100 µL culture from each time point was 10-fold serially diluted for drop plating (10 µL x 3)(94) on TSB blood agar plate for CFU count. Additionally, a 1-mL culture of wild-type strain grown in TSBNH media was collected at day-2, - 3, and -4 to isolate RNA followed by cDNA synthesis using standard protocol. The 1:10 diluted cDNA was used to quantify number of live *P. gingivalis* using 16S rRNA primers specific to *P. gingivalis* (88). The Ct values obtained from unknown samples were used to calculate CFU/mL of *P. gingivalis* from a standard curve prepared with known CFU numbers and their corresponding Ct values (73, 89). The pictures of culture tubes of wild-type strain were also captured to visually represent the growth differences based on the turbidity of bacterial growth in each condition. The assay was repeated at least three times each assay in triplicates. The data is represented as Mean ± SD of three replicates. Students’ two-tailed T-TEST was used for statistical analysis and a p-value of <0.05 (*) or <0.01 (**) was considered statistically significant.

### Atomic force Microscopy (AFM) of *P. gingivalis* wild-type and Δ*gdhA* strains and cell volume calculation

AFM analysis of *P. gingivalis* was carried out following a published protocol(95) with little modification. Wild-type and Δ*gdhA* strains of *P. gingivalis* were grown in TSB or TSBNH media with/out 10 mM Glu-Glu-dipeptideGlu-Glu-dipeptideGlu-Glu-dipeptideto an early exponential phase (OD_600_ = 0.2-0.3). *P. gingivalis* cells from 1 mL culture was then harvested by centrifugation at 4,500 x *g* for 5 min. The cells were then washed twice with 1 mL of sterile (0.1 µm filtered) deionized (DI) water and finally resuspended in 1 mL of DI water. A 50 µL bacterial suspension was then dropped and immobilized on smooth surface of an AFM-grade mica sheet (Electron Microscopy Sciences, # 71855-10) and air dried. Bacterial cells were then fixed with methanol for 2 min (96) and observed under an AFM (Dimension ICON AFM, Bruker, US). The samples were scanned by contact mode using an AFM probe tip (MLCT-BIO-DC, Bruker, US) with a stiffness of 0.07 N/m. The assay was repeated at least three times. The Nanoscope Analysis software v 1.8 (Bruker, US) was used to directly measure the width (W) and length (L) of each bacterium. For the height measurement, the color bar scale for height (Z) in the height sensor image was normalized to minimum data cursor and the section tool was used to measure the height as it was shown before (52, 53). In brief, the XZ data on the graph obtained after cross sectioning an individual bacterium of interest on the image was transferred to excel file and the maximum value of vertical differences between the highest point and base line was used for the height (Z). The cell volume (µm^3^) was calculated using the formula used before,4/3πabc(52, 53)., where a, b, c are the semi axes of solid 3D ellipse and a = L/2, b = W/2, and c = Z/2. A total of at least 25 bacterial cells from each condition were counted. The data is represented as Mean ± SD. Students’ two-tailed T-TEST was used for statistical analysis and a p-value of <0.05 (*) or <0.01 (**) were considered statistically significant.

### Expression and purification of rGdhA (PGN_1367) protein and production of anti-rabbit antibody against rGdhA

The gene for GdhA (PGN_1367) was codon optimized and synthesized before cloning at *Nde*I and *Hind*III sites of pET30a for expression in *E. coli* (GenScript). His tag was placed at the N-terminal of the protein. Ni Column was used for protein purification and the purity was confirmed >90% by SDS-PAGE and specificity of the protein was confirmed by western blotting using mouse anti-His mAb (GenScript # A000186). Excess endotoxin level was removed to keep its level < 1 EU/µg of protein. Purified rGdhA was used to immunize New Zealand Rabbits and antibody titer in serum was checked after 3^rd^ immunization (followed by a booster dose if required) by ELISA (GenScript). The rabbits were sacrificed once the antibody titer was >1:512,000. The antisera were affinity purified against rGdhA. The purified antibody was resuspended in PBS with 0.02% sodium azide and stored at -80°C.

### Organotypic Raft Culture

Once primary GECs and fibroblast cells (FBCs) initially reached 80% confluence, the FBCs were then trypsinized and centrifuged at 59 x g for 5 min. Simultaneously, a collagen raft mixture was prepared on ice via combining 3 mg/ml collagen (Millipore-Sigma, #08-115) with 10X E-Media (DMEM, Millipore-Sigma, #D7777-10L, Ham’s F-12, Millipore Sigma, #N6760) and Reconstitution Buffer (2.2% NaHCO3, 0.15N NaOH, 200 mM HEPES (Millipore-Sigma, #H3375). Trypsin was then decanted, FBS was added to the FBCs at 0.2 mL/raft, the FBCs were added to the collagen raft mixture, and the FBC raft mixture was added to the bottom of 35 mm dishes. The plates were then placed at 37°C in 5% CO2 and were allowed to congeal. Primary GECs were then seeded atop the collagen in KBM media. The rafts were refed daily with KBM for 6 days and were disturbed daily via tapping to prevent attachment. After the sixth day, the rafts were raised via being transferred onto a 50 mm raised wire disk (McMaster Carr, 4195A11) and being placed into a 100 mm dish. After raising, the rafts were refed daily and were permitted to grow for another 4-6 days, pending visible growth of GECs. Rafts were treated with 2 mM EE for 1 h prior to infection with wild-type or Δ*gdhA* strains of *P. gingivalis* for 24 h. Rafts were washed with PBS and placed in 5 mL of 10% Neutral buffered formalin for 18 h prior to being transferred to 70% EtOH until paraffinization and sectioning.

### Immunofluorescence and Confocal Microscopy

To initiate staining, the sectioned organotypic rafts were deparaffinized according to the standard protocol. In brief, sections were heated at 60°C for 30 min, incubated in 100% xylene (Fisher Scientific, #016371.K2) for 7 min, and were dipped 20 times in a secondary 100% xylene solution. The samples then underwent rehydration, where they were dipped 20 times in a series of 100%, 95%, and 70% ethanol solutions before they were placed in PBS for 5 min. Slides were incubated in blocking buffer of 1% BSA in 0.1% triton X-100 in DPBS. Slides were then allowed to partially air-dry prior to incubation with primary antibodies of mouse or rabbit anti-*P. gingivalis* ATCC 33277 (1:200) and rabbit anti-rGdhA (5 µg/mL) or mouse anti-ATG5 (1:200, Proteintech, #66744-1-Ig) antibodies in 1% BSA in PBS for 24 h overnight at 4°C. The samples were then washed briefly in PBS prior to being incubated in corresponding secondary antibodies of Alexa Fluor 488 anti-mouse (1:500, Invitrogen, #A32766) or anti-rabbit (1:500, Invitrogen, #A32731TR) for *P. gingivalis*, Alexa-568 anti-rabbit (1:500, Invitrogen, # A-11011) for GdhA, and anti-mouse 647 (1:500, Invitrogen, #A21235) in 1%BSA in PBS for 1 h at RT. Slides were then dipped into PBS 20 times, after which slides were dipped in 0.1% Sudan Black B (Thermo Scientific, #J62268.22) for 1 h at RT to limit background fluorescence. After 3 times washing in PBS and dehydration by dipping in 70% ethanol, followed by 95% and 100% ethanol, the slides were dipped/incubated in 100% Xylene for 20 times/5 min. The slides were then mounted using Vectashield vibrance antifade mounting medium with DAPI (Vector Laboratories, #H-1800) and glass coverslips.

Imaging was completed the following day using the Leica DM6 CS Stellaris 5 Confocal/Multiphoton System. Control raft sections without either primary antibodies or secondary antibodies were used in the same manner to further verify the specificity of the antibodies used in this study.

### Statistical analysis

All experiments were performed on at least two-three separate occasions with at least two-six technical replicates at each time. The data were initially analyzed by one-way ANOVA and then by Student’s unpaired two-tailed t-test. P-values less than 0.05 (*) or 0.01 (**) determined by the t-test were considered as statistically significant. GraphPad Prism software was used to prepare all the graphs presented in this study.

## Figure Legends for Supplemental figures

**Fig. S1.**
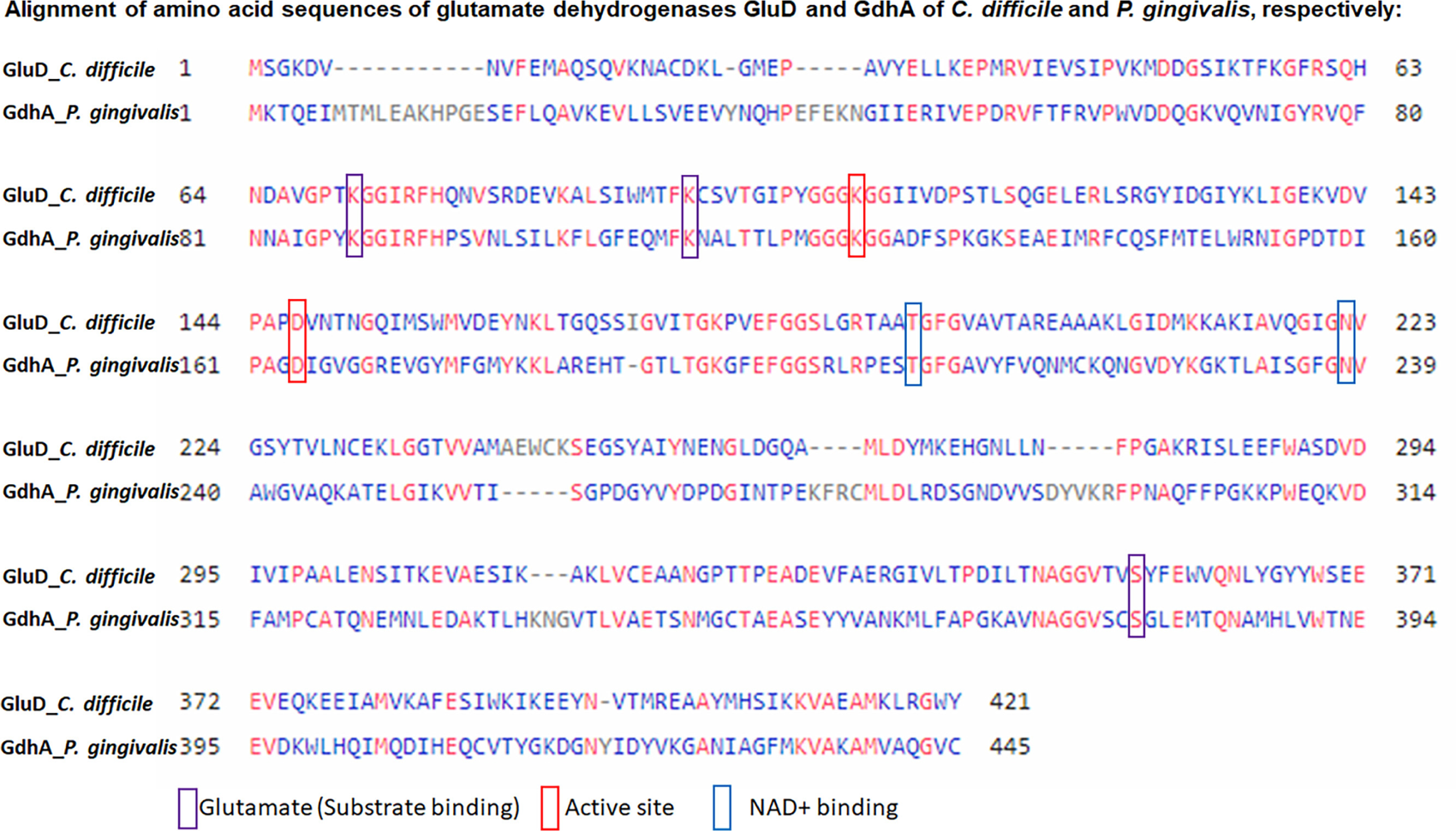
Alignment of amino acid sequences of glutamate dehydrogenase of *C. difficile* and *P. gingivalis*. FASTA sequences of glutamate dehydrogenase Glud (Protein ID# P27346) of *C. difficile* and GdhA or PGN_1367 (protein ID# B2RKJ1) of *P. gingivalis* were aligned using Constraint-based Multiple Alignment Tool (COBALT) (82) available via NCBI BlastP program. The alignment was formatted as compact view using identity conservation setting. Here, red color indicates highly conserved residues and blue indicates less conserved ones. The glutamate (substrate) binding sites, active sites, and the cofactor/coenzyme binding sites are based on the GluD sequences available at www.uniprot.org and highlighted as purple, red, and blue rectangles, respectively. The alignment shows these sites are highly conserved between GluD and GdhA.

## Acknowledgments

This work was primarily supported by funding from the NIH grants R01DE030313, F31DE032273. The authors would like to thank Caleb Wing and Chloe Currie, College of Charleston for their invaluable assistance with the bacterial culturing as volunteer students; MUSC Neuroscience Department for the use of their microscopes; Dr. Shao Yuan of the Hollings Cancer Center Biorepository & Tissue Analysis for his expertise and help with processing of tissue specimens and interpreting them. MUSC Image facilities were supported in part by Hollings Cancer Center Support Grant (P30 CA138313) and the MUSC Digestive Disease Research Cores Center (P30 DK123704).

## References

1. Pelaez-Prestel HF, Sanchez-Trincado JL, Lafuente EM, Reche PA. 2021. Immune Tolerance in the Oral Mucosa. Int J Mol Sci 22.

2. Yang J, Yan H. 2021. Mucosal epithelial cells: the initial sentinels and responders controlling and regulating immune responses to viral infections. Cell Mol Immunol 18:1628–1630.

3. Dale BA. 2002. Periodontal epithelium: a newly recognized role in health and disease. Periodontol 2000 30:70–78.

4. Choi CH, Spooner R, DeGuzman J, Koutouzis T, Ojcius DM, Yilmaz O. 2013. *Porphyromonas gingivalis-*nucleoside-diphosphate-kinase inhibits ATP-induced reactive-oxygen-species via P2X7 receptor/NADPH-oxidase signalling and contributes to persistence. Cell Microbiol 15:961–976.

5. Lee J, Roberts JS, Atanasova KR, Chowdhury N, Yilmaz O. 2018. A novel kinase function of a nucleoside-diphosphate-kinase homologue in *Porphyromonas gingivalis* is critical in subversion of host cell apoptosis by targeting heat-shock protein 27. Cell Microbiol 20:e12825.

6. Lee JS, Chowdhury N, Roberts JS, Yilmaz O. 2020. Host surface ectonucleotidase-CD73 and the opportunistic pathogen, *Porphyromonas gingivalis*, cross-modulation underlies a new homeostatic mechanism for chronic bacterial survival in human epithelial cells. Virulence 11:414–429.

7. Lee K, Roberts JS, Choi CH, Atanasova KR, Yilmaz O. 2018. *Porphyromonas gingivalis* Traffics into Endoplasmic Reticulum-Rich-Autophagosomes for Successful Survival in Human Gingival Epithelial Cells. Virulence doi:10.1080/21505594.2018.1454171:1-26.

8. Roberts JS, Atanasova KR, Lee J, Diamond G, Deguzman J, Hee Choi C, Yilmaz O. 2017. Opportunistic Pathogen *Porphyromonas gingivalis* Modulates Danger Signal ATP-Mediated Antibacterial NOX2 Pathways in Primary Epithelial Cells. Front Cell Infect Microbiol 7:291.

9. Yilmaz O, Verbeke P, Lamont RJ, Ojcius DM. 2006. Intercellular spreading of *Porphyromonas gingivalis* infection in primary gingival epithelial cells. Infect Immun 74:703–710.

10. Yilmaz O, Yao L, Maeda K, Rose TM, Lewis EL, Duman M, Lamont RJ, Ojcius DM. 2008. ATP scavenging by the intracellular pathogen *Porphyromonas gingivalis* inhibits P2X7-mediated host-cell apoptosis. Cell Microbiol 10:863–875.

11. Haffajee AD, Socransky SS. 2005. Microbiology of periodontal diseases: introduction. Periodontol 2000 38:9–12.

12. Patel S, Howard D, Chowdhury N, Derieux C, Wellslager B, Yilmaz O, French L. 2021. Characterization of Human Genes Modulated by *Porphyromonas gingivalis* Highlights the Ribosome, Hypothalamus, and Cholinergic Neurons. Front Immunol 12:646259.

13. Dominy SS, Lynch C, Ermini F, Benedyk M, Marczyk A, Konradi A, Nguyen M, Haditsch U, Raha D, Griffin C, Holsinger LJ, Arastu-Kapur S, Kaba S, Lee A, Ryder MI, Potempa B, Mydel P, Hellvard A, Adamowicz K, Hasturk H, Walker GD, Reynolds EC, Faull RLM, Curtis MA, Dragunow M, Potempa J. 2019. *Porphyromonas gingivalis* in Alzheimer’s disease brains: Evidence for disease causation and treatment with small-molecule inhibitors. Sci Adv 5:eaau3333.

14. Atanasova KR, Yilmaz O. 2014. Looking in the *Porphyromonas gingivalis* cabinet of curiosities: the microbium, the host and cancer association. Mol Oral Microbiol 29:55–66.

15. Liu S, Butler CA, Ayton S, Reynolds EC, Dashper SG. 2024. *Porphyromonas gingivalis* and the pathogenesis of Alzheimer’s disease. Crit Rev Microbiol 50:127–137.

16. Atanasova KR, Yilmaz O. 2015. Prelude to oral microbes and chronic diseases: past, present and future. Microbes Infect 17:473–483.

17. Sheridan M, Chowdhury N, Wellslager B, Oleinik N, Kassir MF, Lee HG, Engevik M, Peterson Y, Pandruvada S, Szulc ZM, Yilmaz O, Ogretmen B. 2024. Opportunistic pathogen *Porphyromonas gingivalis* targets the LC3B-ceramide complex and mediates lethal mitophagy resistance in oral tumors. iScience 27:109860.

18. Holt SC, Ebersole JL. 2005. *Porphyromonas gingivalis*, *Treponema denticola*, and *Tannerella forsythia*: the “red complex”, a prototype polybacterial pathogenic consortium in periodontitis. Periodontol 2000 38:72–122.

19. Socransky SS, Haffajee AD, Cugini MA, Smith C, Kent RL, Jr. 1998. Microbial complexes in subgingival plaque. J Clin Periodontol 25:134–144.

20. Lee JS, Spooner R, Chowdhury N, Pandey V, Wellslager B, Atanasova KR, Evans Z, Yilmaz O. 2020. In Situ Intraepithelial Localizations of Opportunistic Pathogens, Porphyromonas gingivalis and Filifactor alocis, in Human Gingiva. Curr Res Microb Sci 1:7–17.

21. Wisutep P, Kamolvit W, Chongtrakool P, Jitmuang A. 2022. Brain abscess mimicking acute stroke syndrome caused by dual *Filifactor alocis* and *Porphyromonas gingivalis* infections: A case report. Anaerobe 75:102535.

22. Spooner R, Weigel KM, Harrison PL, Lee K, Cangelosi GA, Yilmaz O. 2016. In Situ Anabolic Activity of Periodontal Pathogens *Porphyromonas gingivalis* and Filifactor alocis in Chronic Periodontitis. Sci Rep 6:33638.

23. Lamont RJ, Koo H, Hajishengallis G. 2018. The oral microbiota: dynamic communities and host interactions. Nat Rev Microbiol 16:745–759.

24. Wellslager B, Roberts J, Chowdhury N, Madan L, Orellana E, Yilmaz Ö. 2024. *Porphyromonas gingivalis* activates Heat-Shock-Protein 27 to drive a LC3C-specific probacterial form of select autophagy that is redox sensitive for intracellular bacterial survival in human gingival mucosa. bioRxiv doi:10.1101/2024.07.01.601539:2024.07.01.601539.

25. Dashper SG, Cross KJ, Slakeski N, Lissel P, Aulakh P, Moore C, Reynolds EC. 2004. Hemoglobin hydrolysis and heme acquisition by *Porphyromonas gingivalis*. Oral Microbiol Immunol 19:50–56.

26. Lamont RJ, Jenkinson HF. 1998. Life below the gum line: pathogenic mechanisms of *Porphyromonas gingivalis*. Microbiol Mol Biol Rev 62:1244–1263.

27. Nelson KE, Fleischmann RD, DeBoy RT, Paulsen IT, Fouts DE, Eisen JA, Daugherty SC, Dodson RJ, Durkin AS, Gwinn M, Haft DH, Kolonay JF, Nelson WC, Mason T, Tallon L, Gray J, Granger D, Tettelin H, Dong H, Galvin JL, Duncan MJ, Dewhirst FE, Fraser CM. 2003. Complete genome sequence of the oral pathogenic Bacterium *Porphyromonas gingivalis* strain W83. J Bacteriol 185:5591–5601.

28. Shah HN, Williams RAD. 1987. Utilization of glucose and amino acids by Bacteroides intermedius and Bacteroides gingivalis. Curr Microbiol 15:241–246.

29. Dashper SG, Brownfield L, Slakeski N, Zilm PS, Rogers AH, Reynolds EC. 2001. Sodium ion-driven serine/threonine transport in *Porphyromonas gingivalis*. J Bacteriol 183:4142–4148.

30. Takahashi N, Sato T. 2002. Dipeptide utilization by the periodontal pathogens *Porphyromonas gingivalis*, Prevotella intermedia, Prevotella nigrescens and *Fusobacterium nucleatum*. Oral Microbiol Immunol 17:50–54.

31. Takahashi N, Sato T, Yamada T. 2000. Metabolic pathways for cytotoxic end product formation from glutamate- and aspartate-containing peptides by *Porphyromonas gingivalis*. J Bacteriol 182:4704–4710.

32. Seddon SV, Shah HN, Hardie JM, Robinson JP. 1988. Chemically defined and minimal media forBacteroides gingivalis. Current Microbiology 17:147–149.

33. Wyss C. 1992. Growth of *Porphyromonas gingivalis*, *Treponema denticola, T. pectinovorum*, T. socranskii, and T. vincentii in a chemically defined medium. J Clin Microbiol 30:2225–2229.

34. Masuda K, Tomita K, Hayashi H, Yoshioka M, Hinode D, Nakamura R. 2001. Consumption of Peptide-derived Arginine by a Periodontopathogenic Bacterium, *Porphyromonas gingivalis*. Anaerobe 7:209–217.

35. Spooner R, DeGuzman J, Lee KL, Yilmaz O. 2014. Danger signal adenosine via adenosine 2a receptor stimulates growth of *Porphyromonas gingivalis* in primary gingival epithelial cells. Mol Oral Microbiol 29:67–78.

36. Yoo HC, Yu YC, Sung Y, Han JM. 2020. Glutamine reliance in cell metabolism. Exp Mol Med 52:1496–1516.

37. Yelamanchi SD, Jayaram S, Thomas JK, Gundimeda S, Khan AA, Singhal A, Keshava Prasad TS, Pandey A, Somani BL, Gowda H. 2016. A pathway map of glutamate metabolism. J Cell Commun Signal 10:69–75.

38. Marquez J, de la Oliva AR, Mates JM, Segura JA, Alonso FJ. 2006. Glutaminase: a multifaceted protein not only involved in generating glutamate. Neurochem Int 48:465–71.

39. Mylon E, Heller JH. 1948. Renal glutaminase. Am J Physiol 154:542–548.

40. Errera M. 1949. Liver glutaminases. J Biol Chem 178:483–493.

41. Walker MC, van der Donk WA. 2016. The many roles of glutamate in metabolism. J Ind Microbiol Biotechnol 43:419–430.

42. Mochizuki K. 1999. Purification and characterization of 5-oxo-L-prolinase from Paecilomyces varioti F-1, an ATP-dependent hydrolase active with L-2-oxothiazolidine-4-carboxylic acid. Arch Microbiol 172:182–185.

43. Chen X, Schecter RL, Griffith OW, Hayward MA, Alpert LC, Batist G. 1998. Characterization of 5-oxo-L-prolinase in normal and tumor tissues of humans and rats: a potential new target for biochemical modulation of glutathione. Clin Cancer Res 4:131–138.

44. Ohara-Nemoto Y, Sarwar MT, Shimoyama Y, Kobayakawa T, Nemoto TK. 2020. Preferential dipeptide incorporation of *Porphyromonas gingivalis* mediated by proton-dependent oligopeptide transporter (Pot). FEMS Microbiol Lett 367.

45. Nemoto TK, Ohara Nemoto Y. 2021. Dipeptidyl-peptidases: Key enzymes producing entry forms of extracellular proteins in asaccharolytic periodontopathic bacterium *Porphyromonas gingivalis*. Mol Oral Microbiol 36:145–156.

46. Basic A, Dahlen G. 2023. Microbial metabolites in the pathogenesis of periodontal diseases: a narrative review. Front Oral Health 4:1210200.

47. Girinathan BP, Braun SE, Govind R. 2014. Clostridium difficile glutamate dehydrogenase is a secreted enzyme that confers resistance to H2O2. Microbiology (Reading) 160:47–55.

48. Shetty N, Wren MW, Coen PG. 2011. The role of glutamate dehydrogenase for the detection of *Clostridium difficile* in faecal samples: a meta-analysis. J Hosp Infect 77:1–6.

49. Milner P, Batten JE, Curtis MA. 1996. Development of a simple chemically defined medium for *Porphyromonas gingivalis*: requirement for alpha-ketoglutarate. FEMS Microbiol Lett 140:125–130.

50. Dashper SG, Ang CS, Veith PD, Mitchell HL, Lo AW, Seers CA, Walsh KA, Slakeski N, Chen D, Lissel JP, Butler CA, O’Brien-Simpson NM, Barr IG, Reynolds EC. 2009. Response of *Porphyromonas gingivalis* to heme limitation in continuous culture. J Bacteriol 191:1044–1055.

51. Gallet R, Violle C, Fromin N, Jabbour-Zahab R, Enquist BJ, Lenormand T. 2017. The evolution of bacterial cell size: the internal diffusion-constraint hypothesis. ISME J 11:1559–1568.

52. Malfatti F, Samo TJ, Azam F. 2010. High-resolution imaging of pelagic bacteria by Atomic Force Microscopy and implications for carbon cycling. ISME J 4:427–439.

53. Nishino T, Ikemoto E, Kogure K. 2004. Application of Atomic Force Microscopy to Observation of Marine Bacteria. Journal of Oceanography 60:219–225.

54. Chowdhury N, Norris J, McAlister E, Lau SYK, Thomas GH, Boyd EF. 2012. The VC1777-VC1779 proteins are members of a sialic acid-specific subfamily of TRAP transporters (SiaPQM) and constitute the sole route of sialic acid uptake in the human pathogen Vibrio cholerae. Microbiology (Reading) 158:2158–2167.

55. Lee JS, Yilmaz O. 2021. Key Elements of Gingival Epithelial Homeostasis upon Bacterial Interaction. J Dent Res 100:333–340.

56. Tribble GD, Kerr JE, Wang BY. 2013. Genetic diversity in the oral pathogen *Porphyromonas gingivalis*: molecular mechanisms and biological consequences. Future Microbiol 8:607–620.

57. McDonald ND, Lubin JB, Chowdhury N, Boyd EF. 2016. Host-Derived Sialic Acids Are an Important Nutrient Source Required for Optimal Bacterial Fitness In Vivo. mBio 7:e02237–15.

58. Hamilton PB. 1945. Gasometric determination of glutamine amino acid carboxyl nitrogen in plasma and tissue filtrates by the ninhydrin-carbon dioxide method. Journal of Biological Chemistry 158:375–395.

59. Harris MM. 1943. Studies Regarding a Glutamine-Like Substance in Blood and Spinal Fluid. Science 97:382–383.

60. Atanasova K, Lee J, Roberts J, Lee K, Ojcius DM, Yilmaz O. 2016. Nucleoside-Diphosphate-Kinase of P. gingivalis is Secreted from Epithelial Cells In the Absence of a Leader Sequence Through a Pannexin-1 Interactome. Sci Rep 6:37643.

61. Spooner R, Yilmaz O. 2012. Nucleoside-diphosphate-kinase: a pleiotropic effector in microbial colonization under interdisciplinary characterization. Microbes Infect 14:228–237.

62. Eisele NA, Ruby T, Jacobson A, Manzanillo PS, Cox JS, Lam L, Mukundan L, Chawla A, Monack DM. 2013. Salmonella require the fatty acid regulator PPARdelta for the establishment of a metabolic environment essential for long-term persistence. Cell Host Microbe 14:171–182.

63. Xavier MN, Winter MG, Spees AM, den Hartigh AB, Nguyen K, Roux CM, Silva TM, Atluri VL, Kerrinnes T, Keestra AM, Monack DM, Luciw PA, Eigenheer RA, Baumler AJ, Santos RL, Tsolis RM. 2013. PPARgamma-mediated increase in glucose availability sustains chronic *Brucella abortus* infection in alternatively activated macrophages. Cell Host Microbe 14:159–170.

64. Almeida PE, Carneiro AB, Silva AR, Bozza PT. 2012. PPARgamma Expression and Function in Mycobacterial Infection: Roles in Lipid Metabolism, Immunity, and Bacterial Killing. PPAR Res 2012:383829.

65. Dasgupta S, Rai RC. 2018. PPAR-gamma and Akt regulate GLUT1 and GLUT3 surface localization during Mycobacterium tuberculosis infection. Mol Cell Biochem 440:127–138.

66. Rajaram MV, Brooks MN, Morris JD, Torrelles JB, Azad AK, Schlesinger LS. 2010. Mycobacterium tuberculosis activates human macrophage peroxisome proliferator-activated receptor gamma linking mannose receptor recognition to regulation of immune responses. J Immunol 185:929–942.

67. Abdullah Z, Geiger S, Nino-Castro A, Bottcher JP, Muraliv E, Gaidt M, Schildberg FA, Riethausen K, Flossdorf J, Krebs W, Chakraborty T, Kurts C, Schultze JL, Knolle PA, Klotz L. 2012. Lack of PPARgamma in myeloid cells confers resistance to Listeria monocytogenes infection. PLoS One 7:e37349.

68. Eylert E, Schar J, Mertins S, Stoll R, Bacher A, Goebel W, Eisenreich W. 2008. Carbon metabolism of Listeria monocytogenes growing inside macrophages. Mol Microbiol 69:1008–1017.

69. Madej M, White JBR, Nowakowska Z, Rawson S, Scavenius C, Enghild JJ, Bereta GP, Pothula K, Kleinekathoefer U, Basle A, Ranson NA, Potempa J, van den Berg B. 2020. Structural and functional insights into oligopeptide acquisition by the RagAB transporter from *Porphyromonas gingivalis*. Nat Microbiol 5:1016–1025.

70. Yilmaz O. 2008. The chronicles of *Porphyromonas gingivalis*: the microbium, the human oral epithelium and their interplay. Microbiology (Reading) 154:2897–2903.

71. Syrjanen SM, Alakuijala L, Alakuijala P, Markkanen SO, Markkanen H. 1990. Free amino acid levels in oral fluids of normal subjects and patients with periodontal disease. Arch Oral Biol 35:189–193.

72. Balci N, Kurgan S, Cekici A, Cakir T, Serdar MA. 2021. Free amino acid composition of saliva in patients with healthy periodontium and periodontitis. Clin Oral Investig 25:4175–4183.

73. Lee J, Roberts JS, Atanasova KR, Chowdhury N, Han K, Yilmaz O. 2017. Human Primary Epithelial Cells Acquire an Epithelial-Mesenchymal-Transition Phenotype during Long-Term Infection by the Oral Opportunistic Pathogen, *Porphyromonas gingivalis*. Front Cell Infect Microbiol 7:493.

74. Yilmaz O, Watanabe K, Lamont RJ. 2002. Involvement of integrins in fimbriae-mediated binding and invasion by *Porphyromonas gingivalis*. Cell Microbiol 4:305–314.

75. Oda D, Watson E. 1990. Human oral epithelial cell culture I. Improved conditions for reproducible culture in serum-free medium. In Vitro Cell Dev Biol 26:589–595.

76. Steinhauser C, Heigl U, Tchikov V, Schwarz J, Gutsmann T, Seeger K, Brandenburg J, Fritsch J, Schroeder J, Wiesmuller KH, Rosenkrands I, Walther P, Pott J, Krause E, Ehlers S, Schneider-Brachert W, Schutze S, Reiling N. 2013. Lipid-labeling facilitates a novel magnetic isolation procedure to characterize pathogen-containing phagosomes. Traffic 14:321–336.

77. Roberts-Dalton HD, Cocks A, Falcon-Perez JM, Sayers EJ, Webber JP, Watson P, Clayton A, Jones AT. 2017. Fluorescence labelling of extracellular vesicles using a novel thiol-based strategy for quantitative analysis of cellular delivery and intracellular traffic. Nanoscale 9:13693–13706.

78. Desta IT, Porter KA, Xia B, Kozakov D, Vajda S. 2020. Performance and Its Limits in Rigid Body Protein-Protein Docking. Structure 28:1071–1081 e3.

79. Kozakov D, Beglov D, Bohnuud T, Mottarella SE, Xia B, Hall DR, Vajda S. 2013. How good is automated protein docking? Proteins 81:2159–66.

80. Kozakov D, Hall DR, Xia B, Porter KA, Padhorny D, Yueh C, Beglov D, Vajda S. 2017. The ClusPro web server for protein-protein docking. Nat Protoc 12:255–278.

81. Vajda S, Yueh C, Beglov D, Bohnuud T, Mottarella SE, Xia B, Hall DR, Kozakov D. 2017. New additions to the ClusPro server motivated by CAPRI. Proteins 85:435–444.

82. Papadopoulos JS, Agarwala R. 2007. COBALT: constraint-based alignment tool for multiple protein sequences. Bioinformatics 23:1073–9.

83. Duong-Ly KC, Gabelli SB. 2014. Salting out of proteins using ammonium sulfate precipitation. Methods Enzymol 541:85–94.

84. Nallamsetty S, Waugh DS. 2007. A generic protocol for the expression and purification of recombinant proteins in Escherichia coli using a combinatorial His6-maltose binding protein fusion tag. Nat Protoc 2:383–391.

85. Horton RM, Hunt HD, Ho SN, Pullen JK, Pease LR. 1989. Engineering hybrid genes without the use of restriction enzymes: gene splicing by overlap extension. Gene 77:61–68.

86. Belanger M, Rodrigues P, Progulske-Fox A. 2007. Genetic manipulation of *Porphyromonas gingivalis*. Curr Protoc Microbiol Chapter 13:Unit13C 2.

87. Choi CH, DeGuzman JV, Lamont RJ, Yilmaz O. 2011. Genetic transformation of an obligate anaerobe, P. gingivalis for FMN-green fluorescent protein expression in studying host-microbe interaction. PLoS One 6:e18499.

88. Tran SD, Rudney JD. 1996. Multiplex PCR using conserved and species-specific 16S rRNA gene primers for simultaneous detection of *Actinobacillus actinomycetemcomitans* and *Porphyromonas gingivalis*. J Clin Microbiol 34:2674–2678.

89. Lyons SR, Griffen AL, Leys EJ. 2000. Quantitative real-time PCR for *Porphyromonas gingivalis* and total bacteria. J Clin Microbiol 38:2362–2365.

90. Smalley JW, Silver J, Marsh PJ, Birss AJ. 1998. The periodontopathogen *Porphyromonas gingivalis* binds iron protoporphyrin IX in the mu-oxo dimeric form: an oxidative buffer and possible pathogenic mechanism. Biochem J 331 ( Pt 3):681–685.

91. Okamoto K, Nakayama K, Kadowaki T, Abe N, Ratnayake DB, Yamamoto K. 1998. Involvement of a lysine-specific cysteine proteinase in hemoglobin adsorption and heme accumulation by *Porphyromonas gingivalis*. J Biol Chem 273:21225–21231.

92. Soukos NS, Som S, Abernethy AD, Ruggiero K, Dunham J, Lee C, Doukas AG, Goodson JM. 2005. Phototargeting oral black-pigmented bacteria. Antimicrob Agents Chemother 49:1391–1396.

93. Shah1 HN, Collins2 MD. 1988. Proposal for Reclassification of *Bacteroides asaccharolyticus, Bacteroides gingivalis*, and *Bacteroides endodontalis* in a New Genus, *Porphyromonas*. Int J Syst Evol Bacteriol 38:128–131.

94. Donegan K, Matyac C, Seidler R, Porteous A. 1991. Evaluation of methods for sampling, recovery, and enumeration of bacteria applied to the phylloplane. Appl Environ Microbiol 57:51–56.

95. Allison DP, Sullivan CJ, Mortensen NP, Retterer ST, Doktycz M. 2011. Bacterial immobilization for imaging by atomic force microscopy. J Vis Exp doi:10.3791/2880.

96. Minnerath JM, Roland JM, Rossi LC, Weishalla SR, Wolf MM. 2009. A Comparison of Heat Versus Methanol Fixation for Gram Staining Bacteria. Bioscene 35:36–41.

